# Members of the ELMOD protein family specify formation of distinct aperture domains on the Arabidopsis pollen surface

**DOI:** 10.1101/2021.06.15.448545

**Authors:** Yuan Zhou, Prativa Amom, Sarah H. Reeder, Byung Ha Lee, Adam Helton, Anna A. Dobritsa

## Abstract

Pollen apertures, the characteristic gaps in pollen wall exine, have emerged as a model for studying the formation of distinct plasma-membrane domains. In each species, aperture number, position, and morphology are typically fixed; across species they vary widely. During pollen development certain plasma-membrane domains attract specific proteins and lipids and become protected from exine deposition, developing into apertures. However, how these aperture domains are selected is unknown. Here, we demonstrate that patterns of aperture domains in Arabidopsis are controlled by the members of the ancient ELMOD protein family, which, although important in animals, has not been studied in plants. We show that two members of this family, MACARON (MCR) and ELMOD_A, act upstream of the previously discovered aperture proteins and that their expression levels influence the number of aperture domains that form on the surface of developing pollen grains. We also show that a third ELMOD family member, ELMOD_E, can interfere with MCR and ELMOD_A activities, changing aperture morphology and producing new aperture patterns. Our findings reveal key players controlling early steps in aperture domain formation, identify residues important for their function, and open new avenues for investigating how diversity of aperture patterns in nature is achieved.

## Introduction

As part of cell morphogenesis, cells often form distinct plasma-membrane domains that acquire specific combinations of proteins, lipids, and extracellular materials. Yet how these domains are selected and specified is often unclear. Pollen apertures offer a powerful model for studying this process. Apertures are the characteristic gaps on the pollen surface that receive little to no deposition of the pollen wall exine; during their formation certain regions of the plasma membrane are selected and specified as aperture domains (Zhou and Dobritsa, 2019). Pollen apertures create some of the most recognizable patterns on the pollen surface, usually conserved within a species but highly variable across species (Furness and Rudall, 2004). For instance, in wild-type Arabidopsis pollen, apertures are represented by three long and narrow furrows, equally spaced on the pollen surface and oriented longitudinally (Figure 1A–1A’). In other species, aperture positions, number, and morphologies can be different, suggesting the mechanisms guiding aperture formation are diverse. While the diversity of aperture patterns has captivated scientists for decades (Furness and Rudall, 2004; Matamoro-Vidal et al., 2016; Walker, 1974; Wodehouse, 1935), studies of the associated molecular mechanisms have only recently begun (Dobritsa and Coerper, 2012; Dobritsa et al., 2018; Lee et al., 2018; Reeder et al., 2016; Zhang et al., 2020).

**Figure 1.**
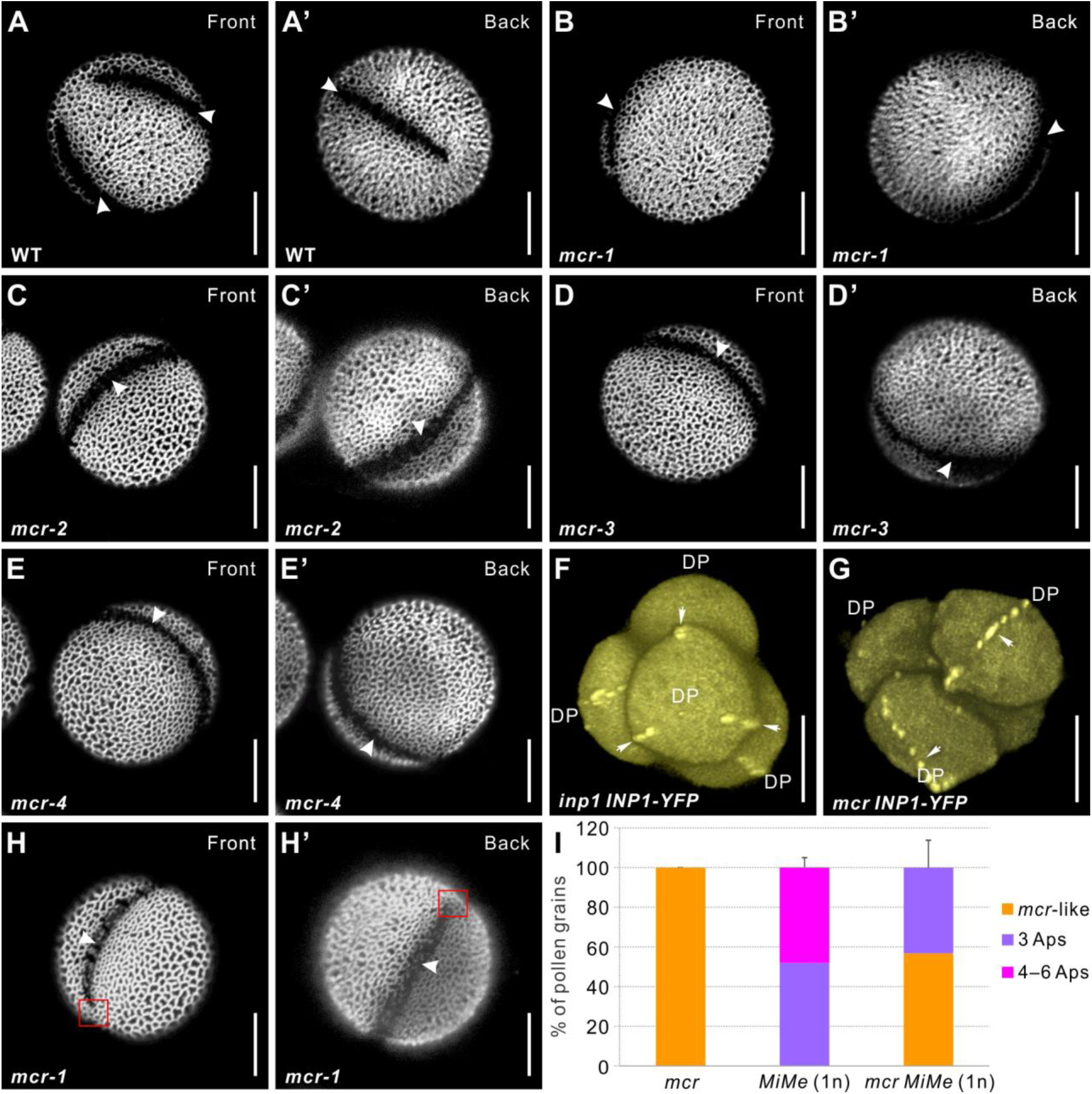
Mutations in *MCR* reduce aperture number. (A–E’) Confocal images of auramine O-stained pollen grains from wild type (L*er*) and four *mcr* EMS mutants. Front (α) and back (α’) show the opposite views of the same pollen grain here and in other figures as indicated. (F–G) 3-D reconstructions of tetrad-stage microspores showing lines of INP1-YFP (arrows) in *inp1* and *mcr* mutants. DP, distal pole. (H–H’) *mcr* pollen with two apertures. Red boxes mark the regions where apertures are not fused. (I) Percentage of pollen grains with indicated number of apertures in pollen populations from *mcr*, 1n *MiMe*, and 1n *mcr MiMe* plants (n = 75–500). Error bars represent SD, calculated from 4–6 independent biological replicates. Apertures are indicated with arrowheads in (A–E’) and (H– H’). Scale bars, 10 μm.

Aperture domains first become visible at the tetrad stage of pollen development, when four sister microspores, the products of meiosis, are held together under the common callose wall and aperture factors, such as INAPERTURATE POLLEN1 (INP1) and D6 PROTEIN KINASE- LIKE3 (D6PKL3) in Arabidopsis and OsINP1 and DEFECTIVE IN APERTURE FORMATION1 (OsDAF1) in rice, accumulate at distinct domains of the microspore plasma membranes (Dobritsa and Coerper, 2012; Dobritsa et al., 2018; Lee et al., 2018; Zhang et al., 2020). These domains become protected from exine deposition and develop into apertures (Dobritsa et al., 2018; Zhang et al., 2020). Yet how aperture domains are selected and what mechanism guides their patterning remains completely unknown.

Recently, we isolated a new Arabidopsis mutant, *macaron* (*mcr*), in which pollen, instead of forming three apertures, develops a single ring-shaped aperture, suggesting that the affected gene is involved in specifying positions and number of aperture domains (Plourde et al., 2019). Here, we perform a detailed analysis of this mutant and identify the *MCR* gene. We demonstrate that it belongs to the ancient family of ELMOD proteins, and that together with another member of this protein family in Arabidopsis, ELMOD_A, MCR acts at the beginning of the aperture formation pathway as a positive regulator of aperture domain specification. We provide evidence that aperture domains are highly sensitive to the levels of MCR and ELMOD_A, which can positively or negatively affect their number. We further demonstrate that a third member of this family, ELMOD_E, has an ability to influence the number, positions, and morphology of aperture domains, and we identify specific protein residues critical for this ability. Our study elucidates key molecular factors controlling aperture patterning and functionally characterizes members of the widespread, yet thus far neglected family of the plant ELMOD proteins.

## Results

### *mcr* mutants develop a single ring-shaped pollen aperture composed of two equidistantly placed longitudinal apertures

In a screen of EMS-mutagenized Arabidopsis plants, we discovered four non-complementing mutants which, instead of three equidistant pollen apertures, produced a single ring-shaped aperture dividing each pollen grain into two equal parts (Figure 1B–1E’). As the mutant phenotype resembled the French meringue dessert, we named these mutations *macaron* (alleles *mcr-1* through *mcr-4*).

Imaging of *mcr* microspore tetrads demonstrated that they develop normally and achieve a regular tetrahedral conformation. The ring-shaped aperture domains in *mcr* microspores, visualized with the help of the reporter INP1-YFP, are positioned so that they pass through the proximal and distal poles of each microspore (Figure 1G; compare with the INP1-YFP localization in the absence of *mcr* mutation in Figure 1F). Thus, like in wild-type pollen, apertures in *mcr* are placed longitudinally. However, while aperture positions in each wild-type microspore are coordinated with aperture positions in its three sisters (Dobritsa et al., 2018; Reeder et al., 2016), in *mcr*, the ring-shaped apertures appear to be placed independently in sister microspores (Figure 1—figure supplement 1). Occasionally, instead of ring-shaped apertures, *mcr* pollen displays two unconnected apertures (Figure 1H–1H’), suggesting that the ring-shaped aperture is a product of a two-aperture fusion. Thus, *mcr* mutations reduce the number of apertures, but do not affect their furrow morphology, longitudinal orientation, and equidistant placement.

### The *mcr* mutation reduces aperture number across different levels of ploidy and arrangements of microspores

We previously showed that aperture number strongly depends on microspore ploidy and is sensitive to cytokinetic defects that disrupt formation of normal tetrahedral tetrads, creating other arrangements of post-meiotic microspores (Reeder et al., 2016). While normal haploid (1n) pollen develops three apertures, diploid (2n) pollen produces either four or a mixture of four and six apertures, depending on whether it was generated through tetrads or dyads. In contrast, 2n *mcr* pollen, produced through either tetrads or dyads, has three equidistant apertures (Plourde et al., 2019), suggesting that the increasing effect of higher ploidy on aperture number is counterbalanced by the defect in the *MCR* function.

We have now extended this analysis by assessing the effects of the *mcr* mutation on aperture formation under additional perturbations of ploidy or post-meiotic microspore arrangement. By creating 1n *Mitosis instead of Meiosis* (*MiMe*) plants (d’Erfurth et al., 2009) with the *mcr* mutation, we generated *mcr* pollen with normal ploidy (1n) via dyads, and not tetrads. As shown previously (Reeder et al., 2016), a majority of the 1n *MiMe* pollen grains (∼60%) develop three normal apertures, with the rest forming mostly six apertures (Figure 1I, Figure 1—figure supplement 2A–C’). Yet, in the pollen of the 1n *mcr MiMe* plants the number of apertures was reduced, with ∼50-70% of pollen developing the *mcr* phenotype (either ring-shaped or two apertures) and the rest forming three apertures (Figure 1I, Figure 1—figure supplement 2D–E’).

We further perturbed microspore formation and ploidy by crossing *mcr-1* with a mutant defective in the *TETRASPORE (TES)* gene. In *tes* mutants, microspore mother cells (MMCs) go through meiosis but fail to undergo cytokinesis, producing large pollen grains with four haploid nuclei and a high number (∼10 or more) of irregularly placed and fused apertures (Reeder et al., 2016; Spielman et al., 1997). Although in the double *mcr tes* mutant apertures are often positioned irregularly and fused together, their number was usually lower (∼4-6) than in the single *tes* mutant (Figure 1—figure supplement 2F–G’). Altogether, these results indicate that *mcr* mutations have an overall reducing effect on aperture number, manifested across different levels of pollen ploidy and post-meiotic microspore arrangements.

### *MCR* acts genetically upstream of the aperture factors *INP1* and *D6PKL3*

In wild-type tetrad-stage microspores, aperture factors INP1 and D6PKL3 localize to the three longitudinal aperture domains of the plasma membrane (Dobritsa and Coerper, 2012; Dobritsa et al., 2018; Lee et al., 2018). Since *mcr* mutation affects INP1-YFP localization, causing it to migrate to a ring-shaped membrane domain (Figure 1G), we tested whether *mcr* also affects the localization of D6PKL3, which likely acts upstream of INP1. We introgressed the previously characterized transgenic reporter *D6PKL3pr:D6PKL3-YFP* (Lee et al., 2018) into the *mcr-1* background. In *mcr* microspores, D6PKL3-YFP re-localized to a single ring-shaped domain (Figure 2A), indicating that MCR acts upstream of both INP1 and D6PKL3.

**Figure 2.**
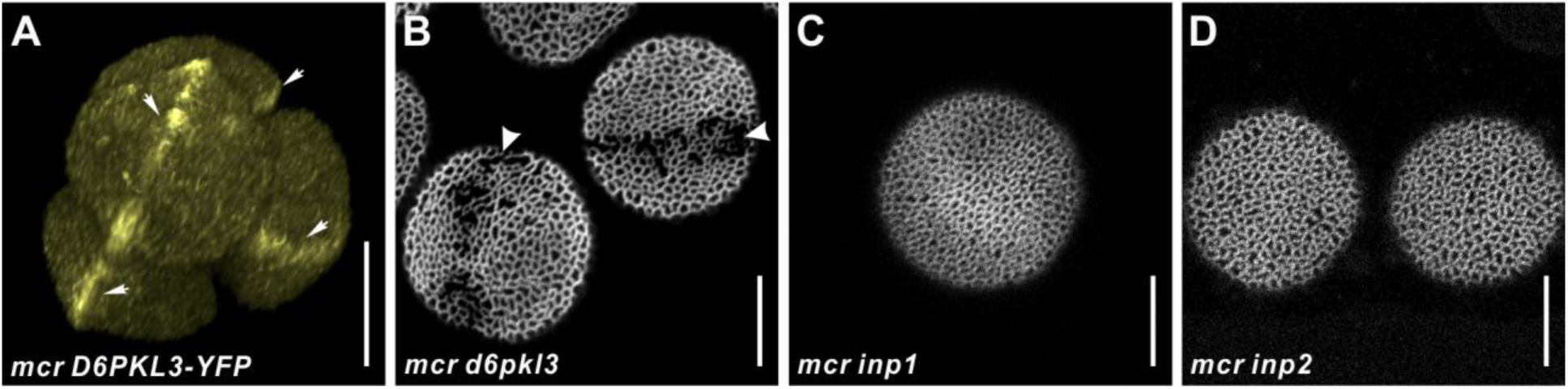
*MCR* acts genetically upstream of the three known aperture factors, *D6PKL3*, *INP1* and *INP2*. (A) 3-D reconstruction of tetrad-stage microspores showing lines of D6PKL3-YFP in *mcr* tetrads. (B–D) Pollen grains of *mcr d6pkl3*, *mcr inp1*, and *mcr inp2* double mutants. Apertures are indicated with arrowheads and D6PKL3-YFP lines are indicated with arrows. Scale bars, 10 μm.

We also examined the genetic interactions between *MCR* and other aperture factors, including the recently discovered *INP2* (Lee et al., 2021), by combining their mutations. *d6pkl3* single mutants develop three apertures partially covered by exine (Lee et al., 2018). Pollen of the *mcr d6pkl3* double mutants developed single ring-shaped apertures that were partially covered by exine, indicating that the two genes have an additive effect on aperture phenotype (Figure 2B). In contrast, pollen grains of *mcr inp1* and *mcr inp2* completely lacked apertures, phenocopying single *inp1* and *inp2* mutants (Dobritsa and Coerper, 2012; Lee et al., 2021) (Figure 2C and 2D). This indicates that *INP1* and *INP2* are epistatic to *MCR*, consistent with their roles of factors absolutely essential for aperture formation.

### MCR is a member of the ancient ELMOD protein family

We mapped the *mcr-1* defect to a 77-kb interval on the second chromosome. One of the 25 genes in this interval, *At2g44770*, had a C-to-T mutation converting a highly conserved Pro165 (see below) into a Ser (Figure 3A, Figure 3—figure supplement 1). Sequencing of *At2g44770* from the other three *mcr* alleles also revealed mutations (Figure 3A, Figure 3—figure supplement 1). *mcr-2* had a G-to-A mutation converting Gly129 into an Asp. *mcr-3* had a G-to-A mutation affecting the last nucleotide of the fifth intron, disrupting the splice acceptor site and causing a frame shift in the middle of the critical catalytic region (see below). In *mcr-4*, no mutations in the coding sequence (CDS) of *At2g44770* were found; however, there was a G-to-A mutation 310 nt downstream of the stop codon in its 3’ untranslated region (3’ UTR), suggesting that the 3’ UTR is important for regulation of this gene (Figure 3A). In addition, plants with T-DNA insertions in this gene (*mcr-5*, *mcr-6*, and *mcr-7*) all produced pollen with the *mcr* phenotype (Figure 3A, Figure 3—figure supplement 2). The T-DNA mutations, however, were hypomorphic, as some pollen with three normal apertures was found in their populations (9% in *mcr-5* (n=179), 13% in *mcr-6* (n=216), and 22% in *mcr-7* (n=78) vs. 0% in *mcr-1* (n=120)).

**Figure 3.**
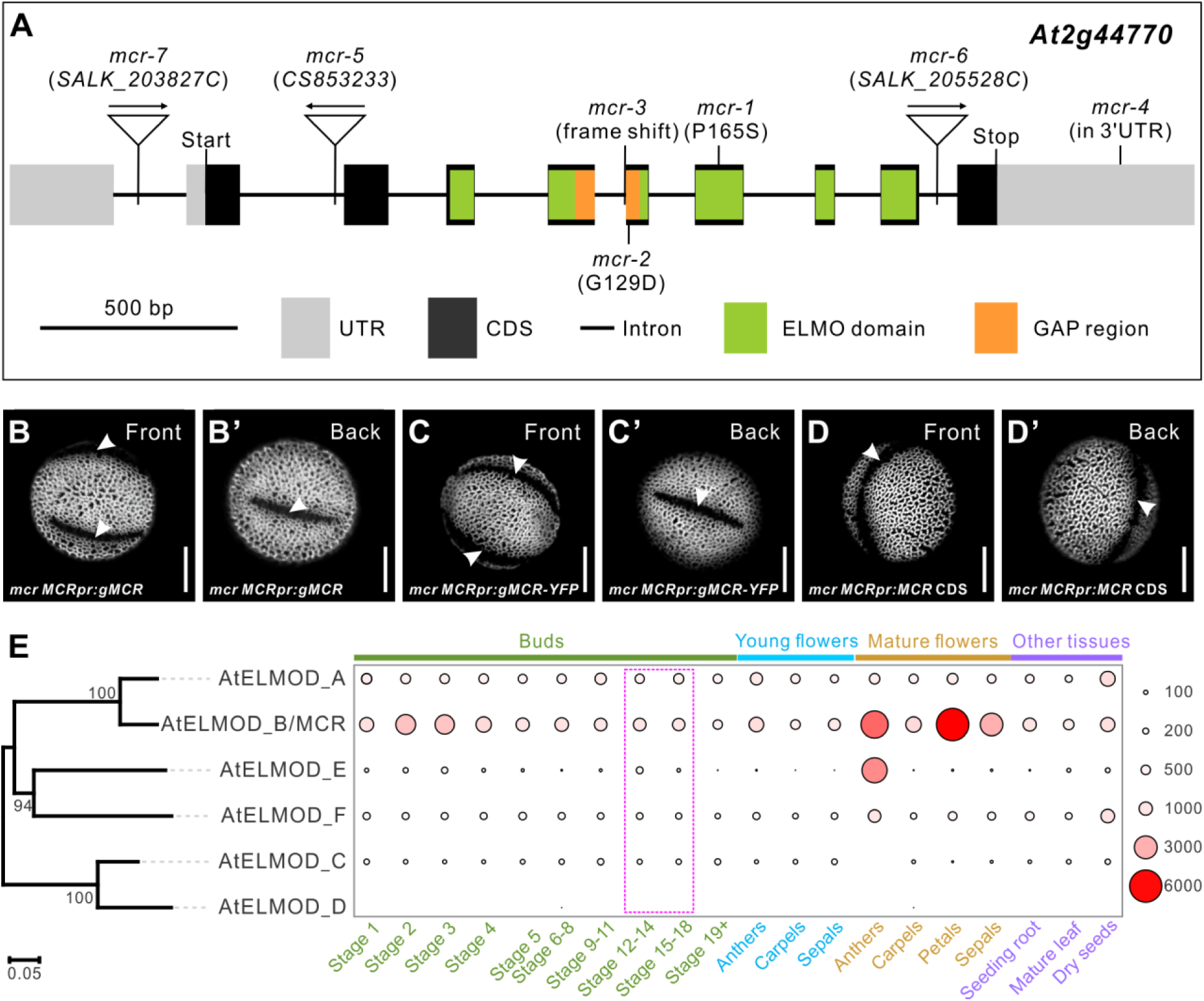
MCR, a member of the ELMOD protein family, is encoded by *At2g44770*. (A) Diagram of the *MCR* gene (*At2g44770*). Positions of seven mutations and several gene and protein regions are indicated. (B–D’) Pollen grains from *mcr* plants expressing *MCRpr:gMCR*, *MCRpr:gMCR-YFP*, and *MCRpr:MCR CDS* constructs. Apertures are indicated with arrowheads. Scale bars, 10 μm. (E) Phylogenetic tree of the Arabidopsis ELMOD proteins and expression patterns of the corresponding genes. Bootstrap values (%) for 1,000 replicates are shown at tree nodes. RNA-seq data obtained from the TRAVA database are presented as a bubble heatmap (values indicate normalized read counts). Magenta box marks the bud stages associated with pollen aperture formation (stages follow the TRAVA nomenclature).

We further verified the identity of *MCR* as *At2g44770* by creating complementation constructs and expressing them in the *mcr-1* mutant. The genomic construct *MCRpr:gMCR* (driven by the 3-kb DNA fragment upstream of the start codon (referred to as the *MCR* promoter) and containing introns and the 0.8-kb region downstream of the stop codon) restored three normal apertures in 10/10 T_1_ transgenic plants (Figure 3B–3B’). A similar genomic construct expressing protein fused at the C-terminus with Yellow Fluorescent protein (YFP) also successfully restored apertures (Figure 3C–3C’). In contrast, the *MCRpr:MCR CDS* construct, which contained only the CDS driven by the *MCR* promoter, did not rescue the *mcr* phenotype (0/6 T_1_ plants had three apertures restored) (Figure 3D–3D’), indicating that additional regulatory regions are required for expression of this gene, consistent with the notion of the 3’ UTR importance. The *MCR* promoter and 3’ UTR were then included in all constructs for which we sought *MCR*-like expression and are herein referred to as the *MCR* regulatory regions.

The protein encoded by *At2g44770* contains the Engulfment and Cell Motility (ELMO) domain (InterPro006816) (Figure 3A, Figure 3—figure supplement 1). In animals, proteins with this domain belong to two families: (1) smaller ELMOD proteins, containing only the ELMO domain and (2) larger ELMO proteins, containing, besides the ELMO domain, several other protein domains (East et al., 2012). The ELMOD family is believed to be the more ancient, with ELMOD proteins already present in the last common ancestor of all eukaryotes, whereas ELMO proteins appeared later in evolution in the opisthokont clade (East et al., 2012). In mammals,

ELMOD proteins act as non-canonical GTPase activating proteins (GAPs) for regulatory GTPases of the ADP-ribosylation factor (Arf) family, a subgroup within the Ras superfamily that includes Arf and Arf-like (Arl) proteins (Bowzard et al., 2007; Ivanova et al., 2014; Turn et al., 2020). Unlike animals, plants only have members of the ELMOD family, and their roles remain essentially uncharacterized.

### Another member of the Arabidopsis ELMOD family, ELMOD_A, is also involved in aperture formation

In Arabidopsis, the ELMOD family consists of six members, ELMOD_A through ELMOD_F (Figure 3E, Figure 3—figure supplement 1), in the nomenclature of (East et al., 2012). MCR is ELMOD_B. One of the other five proteins, ELMOD_A, shares 86% sequence identity with MCR, and the rest have ∼50-55% sequence identity with both MCR and ELMOD_A. Although the ELMOD proteins are broadly expressed in Arabidopsis, young buds at or near the stages when apertures develop express mostly MCR and ELMOD_A (Figure 3E).

Given the high similarity between MCR and ELMOD_A, we wondered if ELMOD_A also aids in aperture formation. We disrupted ELMOD_A with CRISPR/Cas9 (Figure 4A), but it did not affect aperture formation (Figure 4B–4B’). We hypothesized that the lack of phenotype could be due to the ELMOD_A redundancy with MCR. To test this, we crossed the *elmod_a* mutant (carrying the CRISPR/Cas9 transgene) with the *mcr-1* mutant. Already in the F_1_ generation, when all plants were expected to be double heterozygotes, we found several plants producing pollen with the *mcr*-like aperture phenotype (Figure 4C–4C’). Sequencing of the *MCR* and *ELMOD_A* genes from these plants showed that, as expected, they were heterozygous for *MCR*; however, they had homozygous or biallelic mutations in *ELMOD_A*, indicating that the CRISPR/Cas9 transgene continued targeting the wild-type copy of *ELMOD_A* in the F_1_ progeny of the cross.

**Figure 4.**
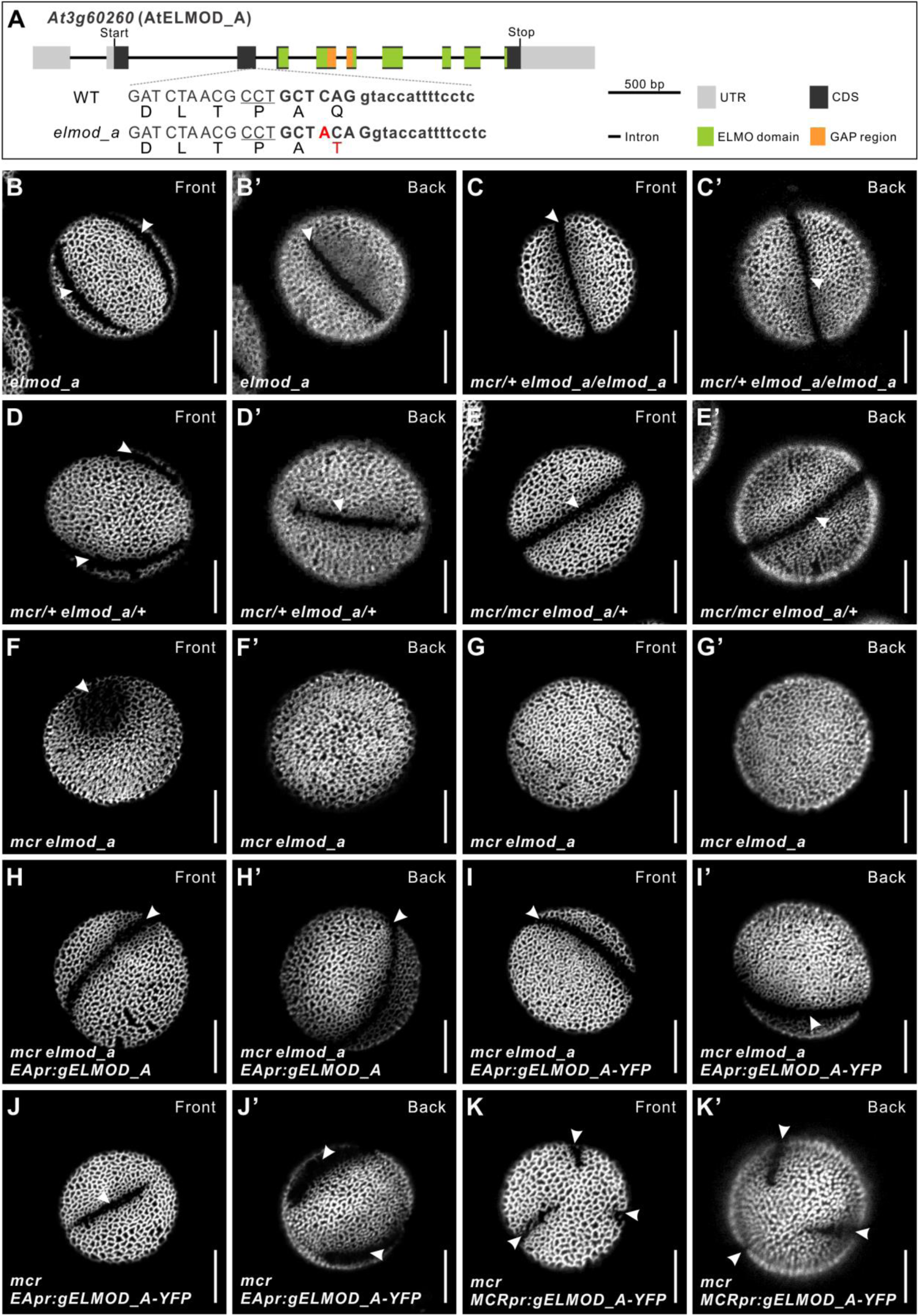
ELMOD_A is involved in aperture formation. (A) Diagram of the *ELMOD_A* gene (*At3g60260*) and the CRISPR/Cas9-induced *elmod_a* mutation. Nucleotide and amino acid changes are indicated with red capital letters. 20-bp target sequence next to the underlined protospacer adjacent motif is shown in bold. Lowercase letters represent sequence of an intron. (B–G’) Pollen grains from *elmod_a* mutant and from the indicated homo- and heterozygous combinations of *elmod_a* and *mcr* mutations. (H–I’) Pollen grains from *mcr elmod_a* plants expressing *EApr:gELMOD_A* and *EApr:gELMOD_A-YFP* constructs. (J–K’) Pollen grains from *mcr* plants expressing *EApr:gELMOD_A-YFP* and *MCRpr:gELMOD_A-YFP* constructs. Apertures are indicated with arrowheads. Scale bars, 10 μm.

The phenotype of these *mcr/+ elmod_a* mutants revealed that in the absence of *ELMOD_A*, *MCR* displays haploinsufficiency. Notably, when at least one wild-type copy of *ELMOD_A* is present, *MCR* is haplosufficient (Figure 4D–4D’). Therefore, these paralogs play redundant roles in the formation of aperture domains. Yet, since MCR can specify three normal apertures in the absence of ELMOD_A but not vice versa, its role appears to be more prominent compared to that of ELMOD_A.

We also tested how the lack of one copy of *ELMOD_A* and both copies of *MCR*, as well as the lack of both genes, would affect aperture formation. In the *mcr elmod_a/+* plants, pollen had the *mcr* phenotype (Figure 4E–4E’). However, when both genes were completely disrupted, the resulting pollen produced either one greatly disrupted aperture with an abnormal, circular morphology and partially covered with exine, or formed no apertures (Figures 4F–4G’). Thus, the simultaneous loss of the two *ELMOD* family genes has a synergistic effect on aperture formation.

To confirm that these defects were caused by mutations in *ELMOD_A* and not off-site CRISPR targeting events, as well as to identify the *ELMOD_A* regulatory regions, we created two *ELMOD_A* genomic constructs driven by the 2-kb region upstream of its start codon – *EApr:gELMOD_A* (which also included a 0.3-kb *ELMOD_A* 3’ UTR) and *EApr:gELMOD_A- YFP* (tagged with YFP and lacking the *ELMOD_A* 3’ UTR) – and transformed them into the *mcr elmod_a* double mutant, which no longer carried the CRISPR/Cas9 transgene. Both constructs successfully rescued formation of apertures (5/5 and 31/33 T_1_ plants, respectively, Figure 4H– 4I’), indicating the selected promoter region is sufficient for *ELMOD_A* functional expression. In addition, when *ELMOD_A* was expressed in the *mcr* single mutant from either its own promoter or from the *MCR* regulatory regions (*MCRpr:gELMOD_A-YFP-MCR3’UTR*), it also complemented the loss of *MCR* (12/12 and 14/14 T_1_ plants) (Figures 4J–4K’).

Thus, both ELMOD_A and MCR participate in aperture domain specification. Formation of three apertures in Arabidopsis pollen requires either two intact copies of *MCR* or at least one copy of each of these two ELMOD family members.

### MCR and ELMOD_A are expressed in the developing pollen lineage but, unlike other aperture factors, do not accumulate at the aperture membrane domains

According to the publicly available RNA-seq data (Klepikova et al., 2016), *MCR* and *ELMOD_A* are both expressed in young buds with pollen at or near the tetrad stage of development (Figure 3E). To confirm that in these buds *MCR* and *ELMOD_A* are expressed in the developing pollen lineage, we created transcriptional reporter constructs *MCRpr:H2B-RFP* and *EApr:H2B-RFP*, expressing the nuclear marker H2B tagged with Red Fluorescent Protein, and transformed them into wild-type Arabidopsis. In the resulting transgenic lines, *MCR* and *ELMOD_A* promoters were active in the developing pollen lineage (MMCs, tetrads, and young free microspores) as well as in somatic anther layers (Figure 5A).

**Figure 5.**
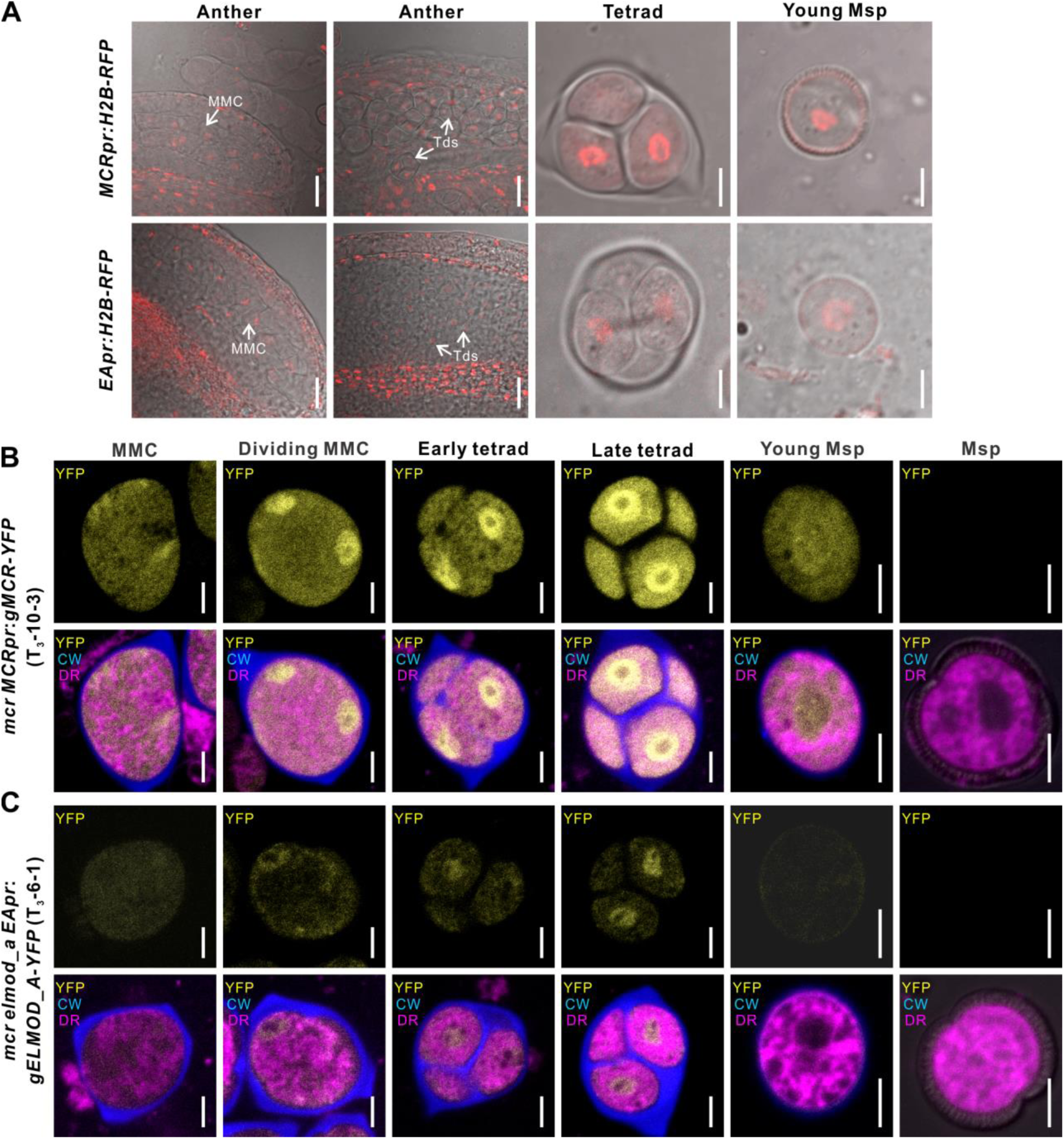
MCR and ELMOD_A do not accumulate at the aperture membrane domains. (A) Confocal images of wild-type anthers, tetrads, and young microspores expressing *MCRpr:H2B-RFP* (upper panels) and *EApr:H2B-RFP* (lower panels). Scale bars, 20 μm for anthers and 5 μm for tetrads and young microspores. (B–C) Confocal images of cells in the developing pollen lineage from *mcr MCRpr:gMCR*-*YFP* (B) and *mcr elmod_a EApr:gELMOD_A*- *YFP* (C) plants. Upper panels: YFP signal. Lower panels: merged signal from YFP (yellow), Calcofluor White (blue, callose wall) and CellMask Deep Red (magenta, membranous structures). Scale bars, 5 μm. Identical staining and color scheme are used for similar images of tetrads in other figures. Abbreviations: CW, Calcofluor White; DR, CellMask Deep Red; MMC, microspore mother cell; Msp, microspore, Td, Tetrad.

To find out if, like the previously discovered aperture factors INP1 and D6PKL3, MCR and ELMOD_A accumulate at the aperture domains of tetrad-stage microspores, we determined the subcellular localization of the YFP-tagged proteins expressed from the translational reporters *MCRpr:gMCR-YFP* and *EApr:gELMOD_A-YFP*, which rescued mutant phenotypes. Consistent with the results from the transcriptional reporters, the YFP signal was present in MMCs, tetrads, and young microspores (Figure 5B–5C). This signal was diffusely localized in the cytoplasm and prominently enriched in the nucleoplasm. No specific enrichment near the plasma membrane was observed. Therefore, MCR and ELMOD_A specify positions and number of aperture domains without visibly congregating there.

### Invariant arginine in the putative GAP region is essential for MCR and ELMOD_A functions

Although ELMOD proteins do not have the typical GAP domain associated with the canonical Arf GAP proteins, they contain a conserved stretch of 26 amino acids, with 13 residues exhibiting a particularly high degree of conservation and forming the consensus sequence WX_3_G(F/W)QX_3_PXTD(F/L)**R**GXGX_3_LX_2_L. In mammalian ELMODs this region is proposed to mediate their Arf/Arl GAP activity (East et al., 2012). The presence of the invariant Arg in this region is of particular importance since the activity of many GAP proteins of the Ras GTPase superfamily, including canonical Arf GAPs, relies on a catalytic Arg (Scheffzek et al., 1998).

Indeed, in mammalian ELMODs, the Arg in this putative GAP region was shown to be essential for their GAP activity, consistent with its role as the catalytic residue (East et al., 2012). Even relatively small changes at this position, such as conversion to Lys, resulted in the complete loss of GAP activity.

Although plant ELMODs have only limited similarity to mammalian proteins (e.g. the Arabidopsis and human ELMODs have ∼20% sequence identity), they contain the same conserved region and invariant Arg residue (Figure 6F). To test if this region is essential for function in MCR and ELMOD_A, we created constructs in which the invariant Arg (R127) was substituted with Lys (*MCRpr:gMCR^R127K^-YFP* and *EApr:gELMOD_A^R127K^-YFP*). These constructs were then expressed, respectively, in the *mcr* and *mcr elmod_a* mutants. Unlike the constructs with the wild-type MCR and ELMOD_A, the R127K constructs, although expressed normally, completely failed to restore the expected aperture patterns (0/8 T_1_ plants for MCR^R127K^; 0/12 T1 plants for ELMOD_A^R127K^), indicating that, like in mammalian ELMODs, the Arg in the putative GAP region is critical for the activity of MCR and ELMOD_A (Figure 6B–6C).

**Figure 6.**
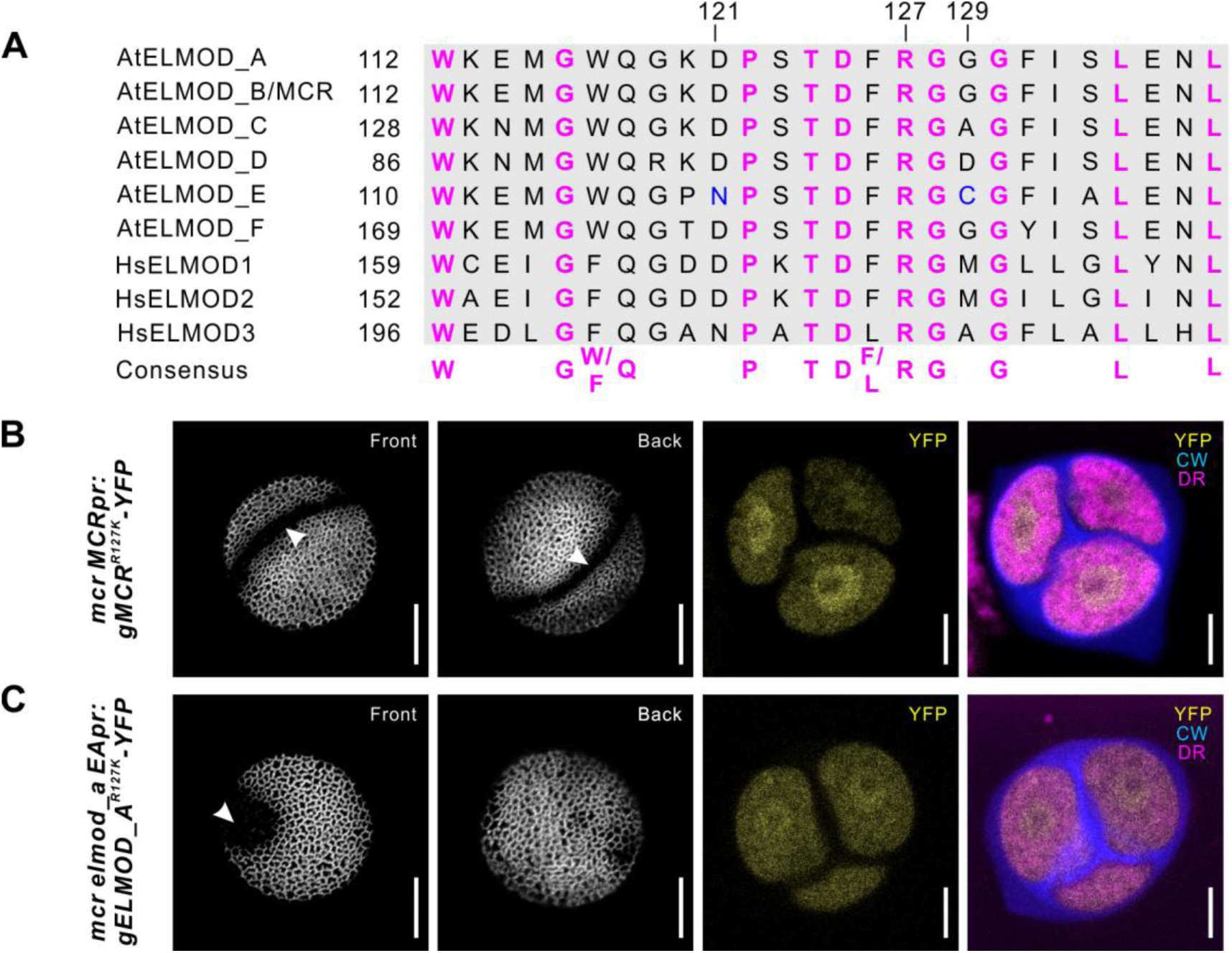
The R127 residue of MCR and ELMOD_A are essential for aperture formation. (A) Sequence alignment of the conserved GAP regions from six Arabidopsis (At) and three human (Hs) ELMOD proteins, along with the consensus sequence. Invariant Arg residue (R127) and two other important residues (121 and 129) are indicated. N121 and C129, essential for AtELMOD_E function, are shown in blue. (B–C) Confocal images of pollen grains and tetrads from *mcr* and *mcr elmod_a* expressing, respectively, *MCRpr:MCR^R127K^-YFP* (B) and *EApr:ELMOD_A^R127K^-YFP* (C). Apertures are indicated with arrowheads. Scale bars, 10 μm for pollen and 5 μm for tetrads.

### The number of developing aperture domains is highly sensitive to the levels of MCR and ELMOD_A

While working with *MCR-YFP* and *ELMOD_A-YFP* transgenic lines, we made a surprising discovery. We noticed that while most of these lines had apertures restored to the expected number (i.e. three apertures for *mcr MCRpr:gMCR-YFP* and a ring-shaped aperture/two apertures for *mcr elmod_a EApr:ELMOD_A-YFP*), in some transgenic T_1_ lines the number of apertures exceeded the expectations: with up to six apertures forming in *mcr MCRpr:gMCR-YFP* and up to four apertures in *mcr elmod_a EApr:gELMOD_A-YFP* (Figure 7A–7B’).

**Figure 7.**
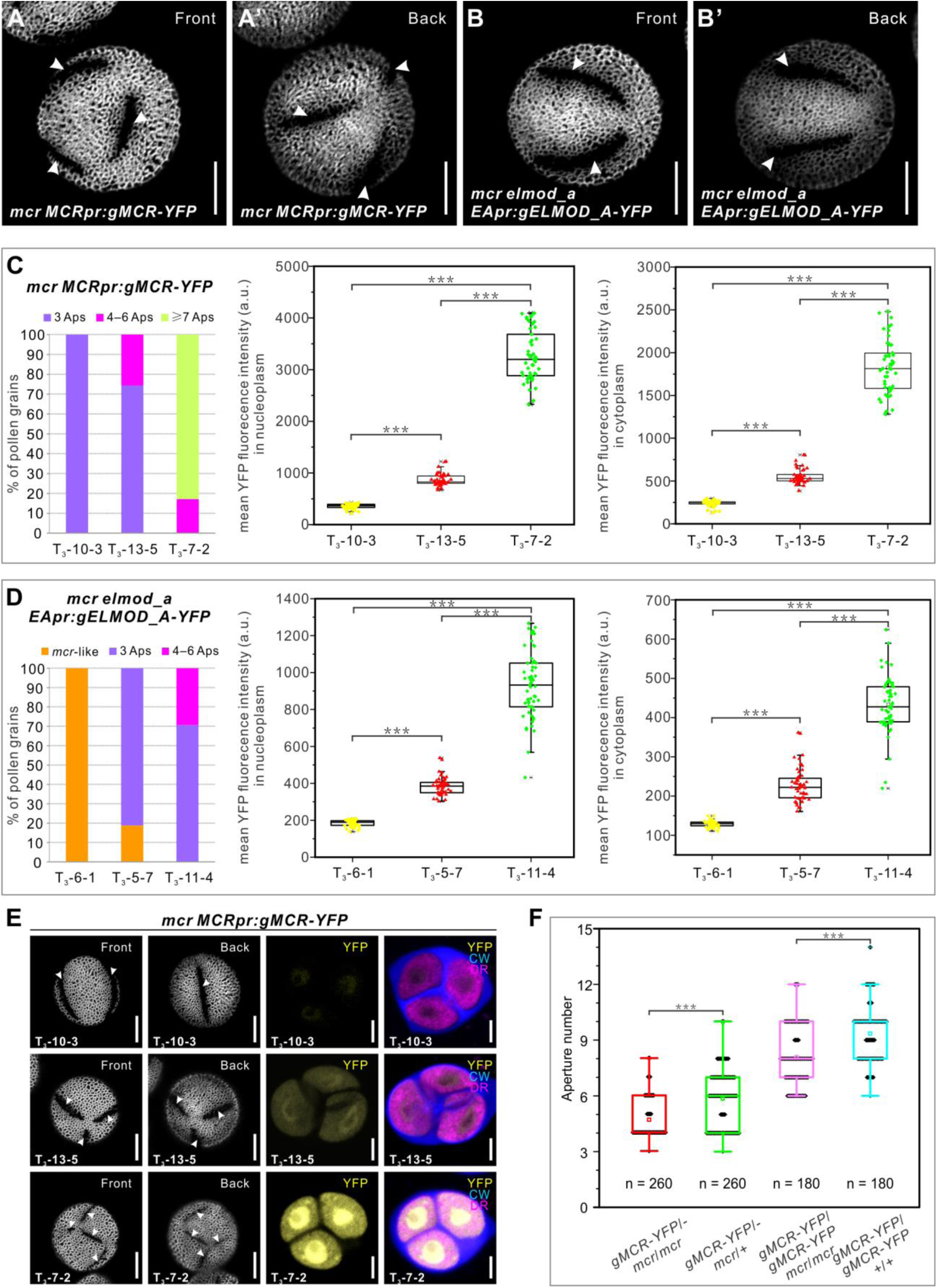
Aperture number is highly sensitive to the levels of MCR and ELMOD_A. (A–B’) Pollen grains from the *mcr MCRpr:gMCR*-*YFP* and *mcr elmod_a EApr:gELMOD_A*-*YFP* transgenic lines, respectively, with six and four apertures. (C–D) Quantification of aperture number and mean YFP signal in three homozygous lines of *mcr MCRpr:*g*MCR*-*YFP* (C) and *mcr elmod_a EApr:*g*ELMOD_A*-*YFP* (D). Stacked bars show the percentage of pollen grains (from ≥3 individual plants) with indicated number of apertures. Boxplots show mean YFP signal in the microspore nucleoplasm and cytoplasm. a. u., arbitrary units. (E) Representative images of pollen grains and tetrads corresponding to data in (C). (F) Boxplots showing aperture number depends on the number of functional copies of *MCR*. Number of analyzed pollen grains (from ≥3 individual plants) is indicated. For all boxplots, boxes represent the first and third quartiles, central lines depict the median, small squares in the boxes indicate the mean values, and small shapes show individual samples. Whiskers extend to minimum and maximum values. \*\*\**p* < 0.001 (two-tailed Student’s t-test). Apertures are indicated with arrowheads. Scale bars, 10 μm for pollen and 5 μm for tetrads.

To test if different aperture numbers could be due to different levels of transgene expression, we examined YFP fluorescence in homozygous lines producing different aperture numbers. For both *MCR* and *ELMOD_A* transgenes, the number of apertures positively correlated with the level of YFP signal in the microspore cytoplasm and nucleoplasm (Figure 7C–7E, Figure 7—figure supplement 1A). In addition, in some lines, the number of apertures further increased in T_2_ or T_3_ generations compared to the numbers in T_1_, consistent with the transgene dosage increasing in later generations due to attaining homozygosity.

To further test the notion that aperture number depends on the *MCR/ELMOD_A* gene dosage/levels of expression, we modulated the dosage of *MCR*, starting with a defined transgene. We crossed a homozygous *mcr MCRpr:gMCR-YFP* plant from line 7-2, commonly producing >6 apertures (Figure 7C), with (1) *mcr* and (2) wild type. In the resulting transgenic F_1_ progeny of the first cross, *MCR* should be expressed from one source – a single copy of the transgene. In the F_1_ progeny of the second cross it should be expressed from two sources – one copy of the transgene plus one of the endogenous gene. In the pollen of these F_1_ plants, the number of apertures correlated with the number of functional copies of *MCR*: pollen of *gMCR-YFP/- mcr/mcr* produced on average 4.68 ± 1.08 apertures compared to 5.85 ± 1.52 apertures in *gMCR- YFP/- mcr/+* (Figure 7F, Figure 7—figure supplement 1B). We further assessed aperture phenotypes in the progeny of these plants that had a homozygous transgene and either zero or two copies of endogenous *MCR*. Both genotypes with the homozygous transgene produced many more apertures compared to plants with the hemizygous transgene, but they also differed significantly from each other, with the number of apertures correlating with the presence of endogenous *MCR* (8.08 ± 1.57 in *MCR-YFP/MCR-YFP mcr/mcr* vs. 9.34 ± 1.50 in *MCR- YFP/MCR-YFP +/+*) (Figure 7F, Figure 7—figure supplement 1B). These results indicate that the process of aperture domain specification is highly sensitive to the levels of MCR and ELMOD_A in developing microspores.

### The ELMOD family in angiosperms has four distinct protein clades, with most species containing two A/B type proteins

To examine the evolutionary history of the plant ELMOD family, we retrieved 561 ELMOD sequences belonging to 178 species across the plant kingdom and used them for a detailed phylogenetic analysis. ELMOD proteins are widespread in plants, suggesting that they perform important functions (Figure 8A).

**Figure 8.**
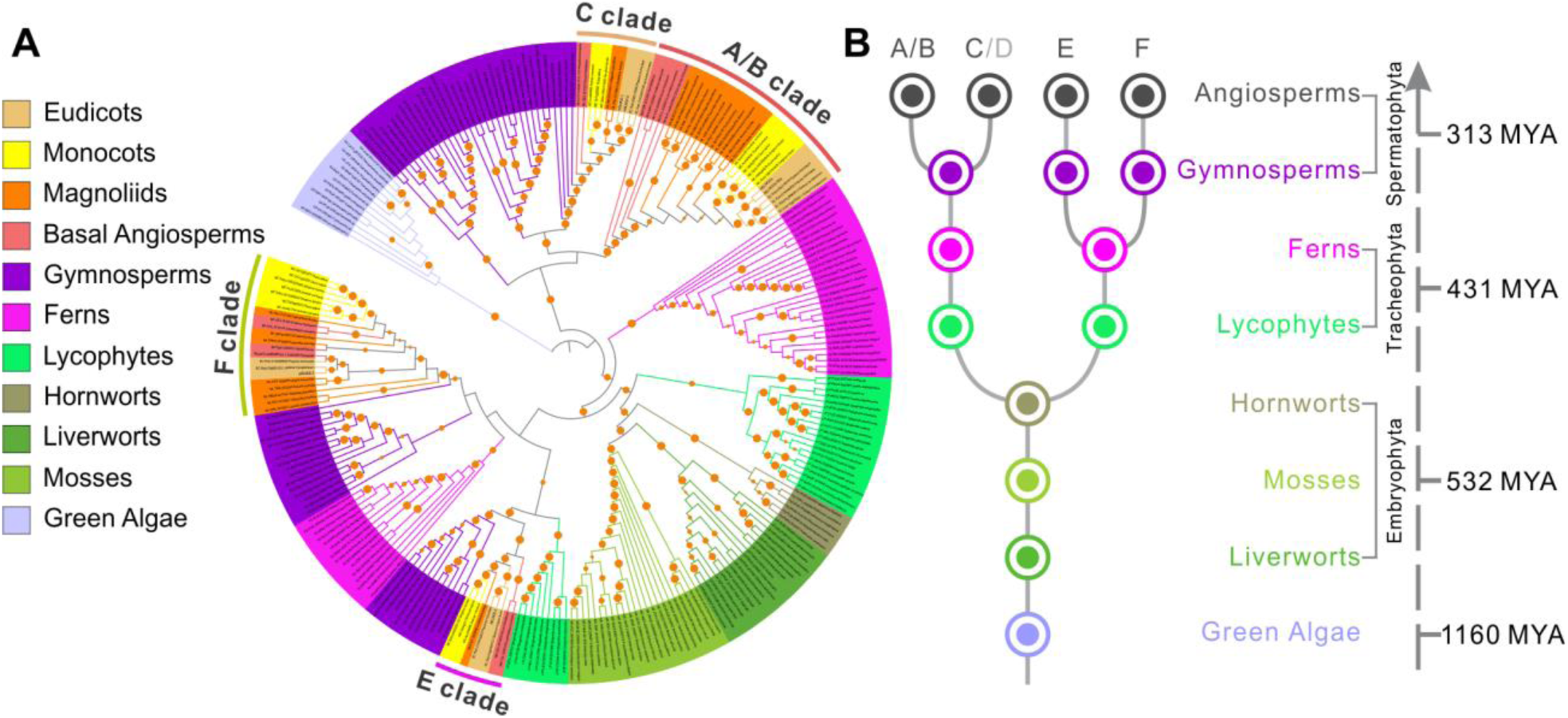
ELMOD proteins exist across the plant kingdom. (A) Maximum likelihood phylogenetic tree of ELMOD proteins across the plant kingdom. The four clades of angiosperm ELMODs are indicated. Orange circles: bootstrap values of 70–100%. (B) Inferred evolutionary history of the *ELMOD* gene family. Dots, inferred ancestral gene number in different plant groups; letters on top, ELMOD clades named after the corresponding Arabidopsis proteins; gray D indicates Arabidopsis *ELMOD_D* is likely a pseudogene; numbers on the right, estimated time of divergence in millions of years (MYA) calculated using the TimeTree database.

Green algae as well as non-vascular land plants (liverworts, mosses, and hornworts) typically have a single ELMOD protein, but an ancestor of lycophytes and ferns had a gene duplication (Figure 8A–8B). Beginning with gymnosperms, the ELMOD family expanded and diversified, with distinct protein groups clustering with the A/B/C clade, the E clade, and the F clade (Arabidopsis proteins were used as landmarks in naming the clades). In early angiosperms, ELMOD proteins separated into four well-supported clades: A/B, C, E, and F (Figure 8A, 8B, Figure 8—figure supplement 1). The split within the aperture factor-containing A/B clade into the separate ELMOD_A and ELMOD_B (MCR) lineages happened late – in the common ancestor of the Brassicaceae family (Figure 8A, Figure 8—figure supplement 1). Yet, in many other angiosperm species, including magnoliids, monocots, basal eudicots, and multiple asterids and rosids, the A/B clade also contains at least two proteins (Figure 8—figure supplement 1). This shows that independent duplications in this lineage happened multiple times, suggesting the existence of strong evolutionary pressure to maintain more than one gene of the A/B type.

### Phylogenetic analysis of the ELMOD family reveals the importance of positions 165 and 129 and suggests ELMOD_D is likely a pseudogene

The extensive number of the retrieved ELMOD sequences allowed us to evaluate conservation of the residues disrupted in MCR by the *mcr-1* and *mcr-2* mutations. Pro165, converted into Ser in *mcr-1* (Figure 3A, Figure 3—figure supplement 1), was present in each of the 553 ELMOD sequences containing this region, suggesting a critical role in protein function. This Pro belongs to the highly conserved WEY**P**FAVAG motif (Figure 3—figure supplement 1) found in all six Arabidopsis ELMODs, as well as in the majority of ELMODs from other plants, including green algae.

The conversion of Gly129 into Asp in *mcr-2* (Figure 3A, Figure 3—figure supplement 1) also represents a very unusual change. Gly129 lies within the putative GAP region, neighboring the critical catalytic Arg127. Notably, except for one likely pseudogene (see below), none of the other 560 retrieved ELMOD sequences has an Asp at that site.

Our careful analysis of residues occupying position 129 in the GAP region across the angiosperm ELMOD proteins led to an interesting discovery. In the 365 analyzed angiosperm sequences, this site is occupied by only three amino acids: Cys, Gly, or Ala. (Earlier diverged plants have Ala or Gly at this site.) Strikingly, we found that all proteins with Cys129 cluster with the E clade, whereas nearly all proteins with Gly129 cluster with either the A/B or the F clades, and nearly all proteins with Ala129 cluster with the C clade. (Only six exceptions were found among the 365 sequences: in five cases, proteins containing Ala129 clustered with the A/B or the F clades, and in one case, a protein with Gly129 clustered with the C clade.) This suggested the intriguing possibility, tested later, that, in angiosperms, residues at position 129 are important for functional differentiation of the ELMOD proteins.

Besides *mcr-2*, the only protein with Asp at position 129 is the Arabidopsis ELMOD_D. However, it has several other features that suggest it is likely a pseudogene. At 213 amino acids, ELMOD_D is markedly shorter than the other five Arabidopsis ELMODs (265 to 323 aa-long): it misses stretches of 52 aa upstream of the GAP region, four aa in the vicinity of the GAP region, and 22 aa at the very C-terminus of the protein (Figure 3—figure supplement 1). It also has major substitutions unique to this protein within or near its GAP region, which change the conserved Gly119 and Leu138 residues into Arg (the numbering within the GAP region is based on the MCR and ELMOD_A sequences) (Figure 3—figure supplement 1). ELMOD_D clusters with the C clade and is most closely related to the Arabidopsis ELMOD_C, indicating it is a product of a very recent duplication (Figure 8—figure supplement 1). While some plants have more than one protein in the C clade, most others, including close relatives of Arabidopsis, have just a single C protein (Figure 8—figure supplement 1), suggesting that a single C-type activity is sufficient for most species. These findings, combined with the extremely low levels of *ELMOD_D* expression in all tissues (Figure 3E), support the hypothesis that this member of the Arabidopsis ELMOD family is likely non-functional.

### ELMOD_E can influence aperture formation and produce a novel aperture pattern

To test if other ELMOD family members, besides MCR and ELMOD_A, might be involved in the aperture formation, we first examined the phenotypes of *elmod_c*, *elmod_d*, *elmod_e* and *elmod_f* single mutants carrying T-DNA insertions in their CDS (Figure 9—figure supplement 1A). All mutants displayed normal aperture patterns (Figure 9—figure supplement 1B–1E’). For *ELMOD_E*, we also created a mutant allele by CRISPR/Cas9, generating an early frame shift (Figure 9—figure supplement 1A), which also produced three normal apertures (Figure 9— figure supplement 1F–1F’). Additionally, we combined mutations in these four genes with *mcr*, creating a series of double mutants, which exhibited *mcr* phenotypes (Figure 9—figure supplement 1G–J’), indicating that, unlike *ELMOD_A*, the other four *ELMOD* genes do not interact synergistically with *MCR* in aperture formation.

**Figure 9.**
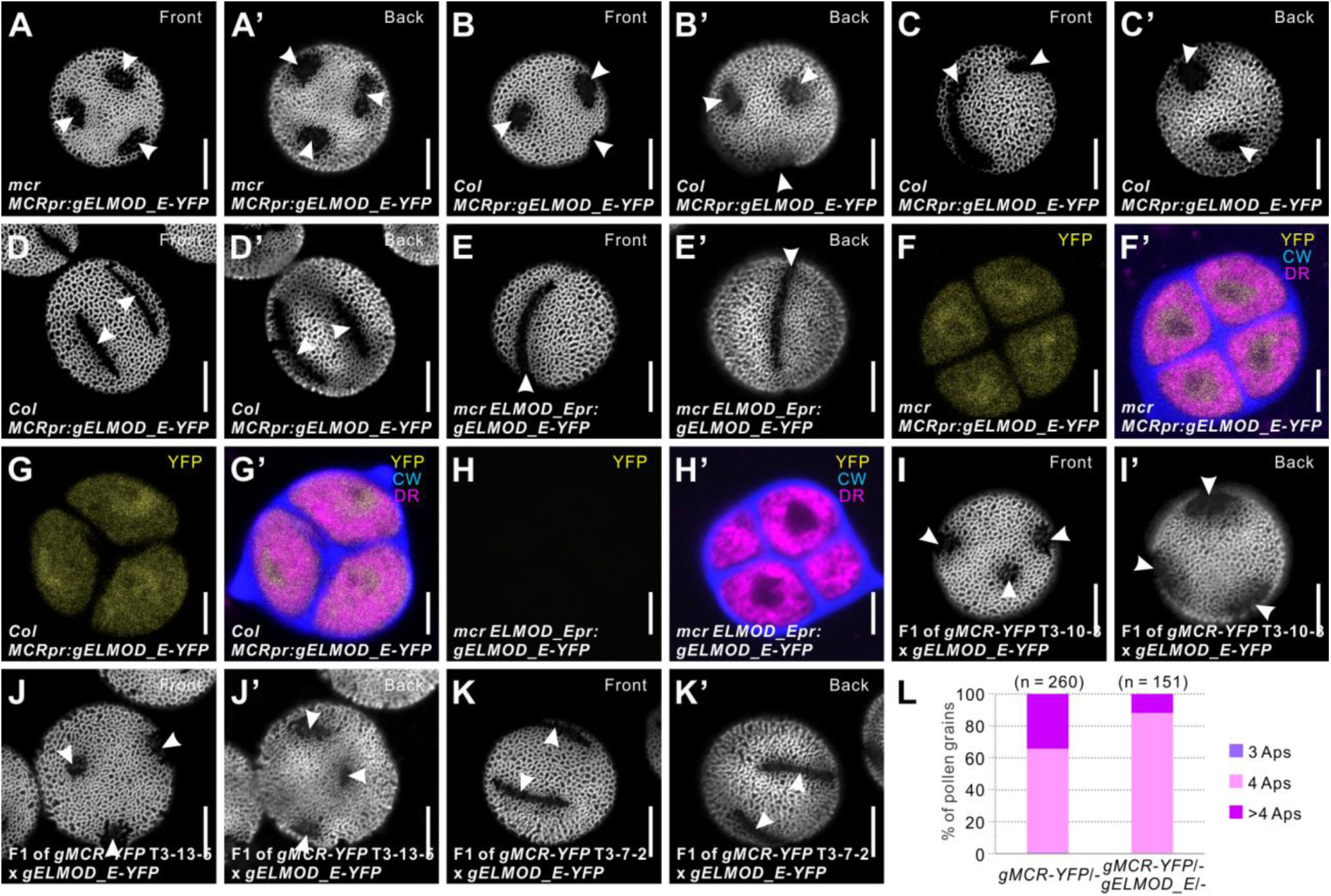
Arabidopsis ELMOD_E can affect aperture patterns. (A–D’) Pollen grains from *mcr* (A–A’) and Col-0 (B–D’) plants expressing *MCRpr:gELMOD_E-YFP*. (E–E’) Pollen grain from *mcr* plants expressing *ELMOD_Epr:gELMOD_E-YFP*. (F–H’) Confocal images of tetrads expressing *MCRpr:gELMOD_E-YFP* and *ELMOD_Epr:gELMOD_E-YFP*. Adjacent panels show YFP signal (α) and merged signal (α’) from YFP, Calcofluor White (CW), and CellMask Deep Red (DR). (I–K’) Pollen grains from the F_1_ plants produced by crossing *mcr MCRpr:gELMOD_E-YFP* with three T_3_ lines of *mcr MCRpr:gMCR-YFP* (with single homozygous insertions of the *MCR*-*YFP* transgene, expressed, respectively, at low, medium, and high levels). (L) Percentage of pollen grains with indicated number of apertures in the pollen populations from F_1_ progeny of the *mcr MCRpr:gMCR-YFP* T_3_-7-2 line crossed with *mcr* or with *mcr MCRpr:gELMOD_E-YFP.* Number of analyzed pollen grains (from at least two individual plants) is indicated. Apertures are indicated with arrowheads. Scale bars, 10 μm for pollen and 5 μm for tetrads.

We also assessed the ability of these four genes to rescue the *mcr* aperture phenotype by expressing them from the *MCR* regulatory regions. The expression of the *ELMOD_D* and *ELMOD_F* did not produce changes in the aperture pattern (6/6 and 22/22 T_1_ plants) (Figure 9— figure supplement 2A–2B’). As all the transgenes contained YFP, we monitored their expression. While the ELMOD_F-YFP was expressed at a level comparable with that of the MCR-YFP and ELMOD_A-YFP transgenes, ELMOD_D-YFP was expressed at a very low level (Figure 9— figure supplement 2C–2D’), consistent with the hypothesis that *ELMOD_D* is a pseudogene.

Since the *ELMOD_D* transgene was expressed from the *MCR* regulatory regions, its expression might be regulated at the post-transcriptional level. Unlike *ELMOD_D* and *ELMOD_F*, *ELMOD_C* had some ability, albeit limited, to restore three apertures in *mcr* (Figure 9—figure supplement 2E–2H’).

Most interestingly, the expression of *ELMOD*_*E* in *mcr* led to a neomorphic phenotype: instead of narrow longitudinal furrows, pollen of all seven T_1_ plants developed multiple short, round apertures (Figure 9A–9A’). To see if this effect was limited to the *mcr* background, we transformed the *MCRpr:ELMOD_E-YFP* construct into wild-type Col-0 plants. The T_1_ plants showed a range of aperture phenotypes (Figure 9B–9D’), yet multiple round apertures were commonly present, suggesting that *ELMOD*_*E* exerts a dominant negative effect when misexpressed in developing microspores.

We then tested whether *ELMOD_E* would have the same effect on aperture patterns when expressed from its own promoter. However, none of the 12 T_1_ transgenic *mcr* plants expressing the *ELMOD_Epr:ELMOD_E-YFP* construct had any changes in the *mcr* aperture phenotype (Figure 9E–9E’). Analysis of the YFP signal showed that this gene is expressed in tetrad-stage microspores at much lower levels from its own promoter than from the *MCR* promoter (Figure 9F–9H’). Thus, while ELMOD_E can influence aperture patterns, it is likely not normally involved in this process in Arabidopsis.

To test if differences in the MCR levels could impact the ability of transgenic ELMOD_E to produce round apertures, we crossed a *mcr MCRpr:gELMOD_E-YFP* line with the above described *mcr MCRpr:gMCR-YFP* lines 10-3, 13-5, and 7-2 that express MCR, respectively, at low, medium, and high levels (Figure 7C). In the F_1_ progeny of crosses with lines 10-3 and 13-5, pollen still produced round apertures (Figure 9I–9J’). Yet, in the F_1_ progeny of the cross with line 7-2 expressing MCR at high level, furrow aperture morphology was restored (Figure 9K– 9K’), suggesting that high level of MCR can counteract the neomorphic activity of ELMOD_E. The number of furrows produced by the F_1_ progeny of that cross was lower than in the F_1_ progeny of the cross between line 7-2 and *mcr*, which had the same *MCR* dosage but lacked the *ELMOD_E* transgene (Figure 9L). These data support the idea that MCR and ELMOD_E likely compete for the same interactors.

### Residues 121 and 129 in the GAP region are important for the MCR- and ELMOD_E- specific functions in aperture formation

The different aperture phenotypes of *mcr MCRpr:gMCR-YFP* and *mcr MCRpr:gELMOD_E- YFP* lines gave us an opportunity to test the hypothesis that residues at position 129 are important for functional differentiation of ELMODs from different clades. For E-clade proteins, we also noticed that Cys129 was always found together with Asn121. These residues are unique to this clade: 100% of the E-clade sequences (n=69) have Asn121/Cys129 vs. 0% of sequences from the other clades (n=297). Thus, this combination could be important for the E-clade functions. In the other three clades, position 121 is always occupied by Asp.

To investigate the importance of sites 121 and 129 for MCR and ELMOD_E functions, we created six constructs in which one or both residues at these positions were replaced with the residues typical of the other clade and expressed them in the *mcr* mutant. The MCR proteins carrying the E-specific residues at both positions (MCR^D121N/G129C^) or at the position 121 (MCR^D121N^) still retained most of the MCR function, with most T_1_ plants producing three or more furrow apertures in most of their pollen grains (7/9 and 12/12 T_1_ plants, respectively) (Figure 10A, 10B, Figure 10—figure supplement 1A). However, when the E-specific residue was present only at position 129 (MCR^G129C^), MCR protein became less active, with only 5 out of 11 T_1_ plants producing three furrow apertures in all or most of their pollen (Figure 10C). In the rest of these T_1_ plants, the *mcr* phenotype was not rescued or was rescued poorly, with <30% of pollen grains forming three apertures.

**Figure 10.**
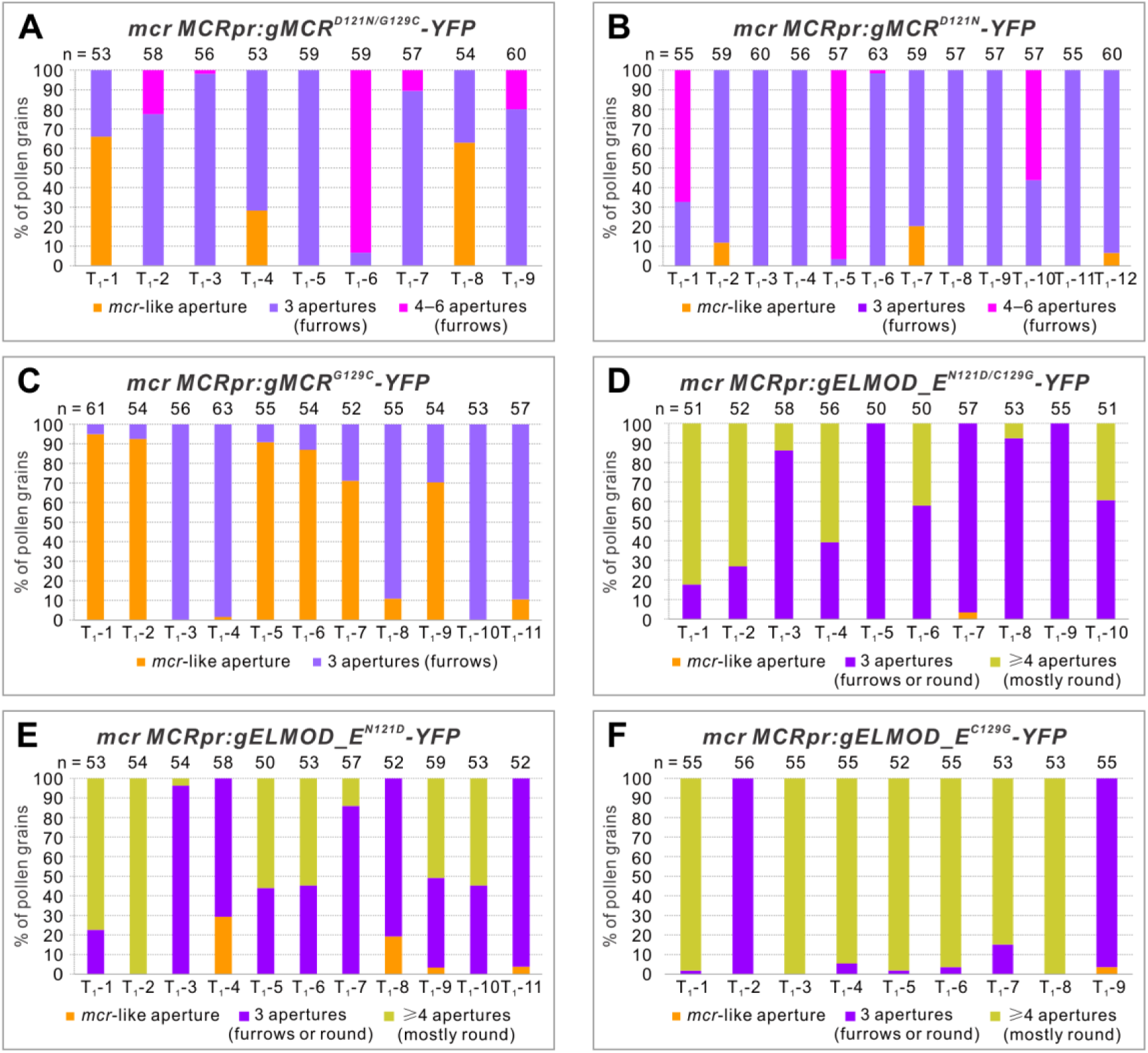
Residues 121 and 129 in the GAP region are important for MCR- and ELMOD_E-specific functions in aperture formation. Percentage of pollen grains with indicated number of apertures in the pollen populations from independent T_1_ *mcr* plants expressing variants of *MCRpr:gMCR-YFP* (A–C) or *MCRpr:ELMOD_A-YFP* (D–F) with residues 121 and/or 129 mutated. Number of analyzed pollen grains is indicated above the bars.

Experiments with the ELMOD_E proteins carrying the MCR residues at positions 121 and 129 confirmed the importance of Asn121 and Cys129 for the ELMOD_E neomorphic activity. In the case where both residues were replaced with the MCR residues (ELMOD_E^N121D/C129G^), ELMOD_E largely lost its ability to create round apertures and instead often restored three furrow-like apertures, thus acting like MCR (Figure 10D, Figure 10—figure supplement 1B). In the cases when only one residue was changed (ELMOD_E^N121D^ and ELMOD_E^C129G^), the mutant ELMOD_E proteins were still often able to produce multiple round apertures, although three normal furrows or a mixture of furrows and round apertures were also produced, suggesting that the single mutations reduced the ELMOD_E activity, but not eliminated it entirely (Figure 10E, 10F, Figure 10—figure supplement 1B).

Taken together, these results show that residues at positions 121 and 129 in the GAP region provide important contributions to the specific function of each protein. Yet they are less critical for MCR, in accord with the fact that Asp121 and Gly129 are not unique to the A/B clade. In the case of ELMOD_E, the E-clade-specific combination of Asn121/Cys129 appears to be essential for its distinct activity. When both residues undergo MCR-like changes, ELMOD_E loses its neomorphic activity, instead becoming capable of carrying out the MCR role in aperture formation.

## Discussion

How developing pollen grains create beautiful and diverse geometrical aperture patterns has been a long-standing problem in plant biology (Fischer, 1889; Ressayre et al., 2002; Wodehouse, 1935). In this study, we uncovered the first set of molecular factors, belonging to the ELMOD protein family, that have a clear ability to regulate the number, positions, and morphology of aperture domains. MCR and its close paralog ELMOD_A act as (somewhat) redundant positive regulators of furrow aperture formation in Arabidopsis.

Our genetic analysis places MCR and ELMOD_A at the beginning of the aperture formation pathway, upstream of the previously discovered aperture factors D6PKL3, INP1, and, likely, INP2, the recently identified partner of INP1. Previous studies showed that INP1 and INP2 act as the executors of the aperture formation program, absolutely essential for aperture development but not able on their own to influence the number and positions of aperture domains (Dobritsa et al., 2018; Lee et al., 2021; Li et al., 2018; Reeder et al., 2016). D6PKL3 was proposed to act upstream of INP1, defining the features of aperture domains, yet it also largely lacks the ability to initiate completely new domains (Lee et al., 2018; Zhou and Dobritsa, 2019). In *mcr* microspores, D6PKL3 and INP1 re-localize to the ring-shaped aperture domains (Figure 1G and 2A), indicating that they become attracted to the newly specified aperture domains and the ELMOD proteins act as patterning factors, contributing to symmetry breaking and selection of sites for aperture domains.

Our data demonstrate that the aperture domains forming in each microspore are highly sensitive to the ELMOD_A/MCR protein dosage (Figure 4C–4G’, Figure 7, Figure 7—figure supplement 1). Increased dosage leads to a higher number of apertures, while decreased dosage results in fewer. Thus, modulation of ELMOD protein levels offers a mechanism for creating different aperture patterns in different species. Interestingly, within the genus *Pedicularis,* some species display the *mcr*-like ring-shaped apertures, while others produce three apertures (Wang et al., 2009, 2017), which could be due to variations in ELMOD proteins or their effectors or regulators. Importantly, while great diversity of pollen aperture numbers is found across plant species, within a species, this trait tends to be very robust. For example, in wild-type Arabidopsis, the number of apertures rarely deviates from three (Reeder et al., 2016). Our results, therefore, imply that, usually, levels of MCR and ELMOD_A are likely very tightly controlled. Our transgenic constructs likely miss some regulatory elements controlling expression of these genes from the endogenous sites in the genome. Additionally, there might be other mechanisms to precisely regulate the activity of these *ELMOD* genes.

The discovery that *ELMOD_E* can also influence aperture patterns in Arabidopsis and create multiple round apertures instead of three furrows (Figure 9A–9B’) suggests that the regulation of *ELMOD_E* might also contribute to the diversity of aperture patterns in nature. In Arabidopsis, *ELMOD_E* does not seem to be usually involved in aperture formation. Yet, when misexpressed from the *MCR* regulatory regions, it interferes with MCR and ELMOD_A activity (Figure 9I– 9L), resulting in the formation of new aperture domains.

ELMODs are ancient proteins, predicted to have been present in the last common ancestor of all eukaryotes (East et al., 2012). In animals, these proteins act as non-canonical GTPase activating proteins (GAPs), regulating activities of both Arf and Arl GTPases (Bowzard et al., 2007; Ivanova et al., 2014; Turn et al., 2020). Arf GTPases are commonly associated with the recruitment of vesicle coat proteins to different membrane compartments to initiate vesicle budding and trafficking, while the roles of the related Arl proteins are less understood and likely more diverse (Sztul et al., 2019). Although the function of ELMOD proteins in plants is unknown, their presence in green algae and other basal plants suggests that they have been playing important roles in plant cells since their inception. Our phylogenetic analysis indicates that this family in plants is monophyletic, and the genes have duplicated and diversified over the course of plant evolution.

The angiosperm ELMOD family has four distinct clades (Figure 8A, 8B, Figure 8—figure supplement 1). In many species, the A/B clade, containing MCR and ELMOD_A, has two or more proteins, due to independent duplications that occurred multiple times in evolution. This suggests that species might be under a selective pressure to keep more than one A/B type protein, implying that the processes in which these proteins are involved (e.g. aperture formation) benefit from genetic redundancy and, thus, are highly important.

Further studies will be required to establish the biochemical role of plant ELMOD proteins. Like their animal counterparts, plant ELMODs may be involved in regulation of Arf/Arl activities. The protein region proposed to be the GAP region in mammalian ELMODs (East et al., 2012) is conserved in plant proteins, and the invariant Arg residue believed to be catalytic in mammalian ELMODs is also necessary for function in MCR and ELMOD_A (Figure 6B–6C). Interestingly, some positions within the conserved GAP region show strict residue specificity in different clades, suggesting they could be important for functional diversity of these proteins. Consistent with this, we found the combination of Asn121/Cys129 to be key for the ELMOD_E neomorphic aperture-forming activity (Figure 10D–10F).

Arabidopsis has 19 ARFs and ARLs, and with few exceptions, most remain uncharacterized (Delgadillo et al., 2020; Gebbie et al., 2005; McElver et al., 2000; Singh et al., 2018; Vernoud et al., 2003; Xu and Scheres, 2005). The roles attributed to members of this family – e.g. in secretion, endocytosis, activation of phosphatidyl inositol kinases, and actin polymerization (Singh and Jürgens, 2018; Sztul et al., 2019) – are all potentially fitting with the formation of distinct aperture domains. If ELMODs are indeed the negative regulators of ARF/ARL GTPases, this would suggest that activity of these GTPases can inhibit formation of aperture domains and they have to be kept in check.

Alternatively, plant ELMODs could have evolved new functions. Interestingly, the only study done so far on an ELMOD protein in plants (Hoefle and Hückelhoven, 2014) pulled out the barley homolog of ELMOD_C in a yeast two-hybrid screen as an interactor of a ROP GAP, a GAP for a different class of small GTPases, Rho-of-plants. Rho GTPases (including ROPs) are well-known regulators of cell polarity and domain formation (Feiguelman et al., 2018; Yang and Lavagi, 2012), so their involvement in aperture formation cannot be excluded.

In summary, we presented critical players in the process of patterning the pollen surface. These players belong to the ELMOD protein family, which, while undoubtedly important, has not yet been characterized in plants. Future studies should focus on identifying the interactors of the ELMOD proteins and on understanding the mechanisms through which they specify positions and shape of aperture domains without noticeably accumulating at these regions.

## Materials and methods

### Key resources table

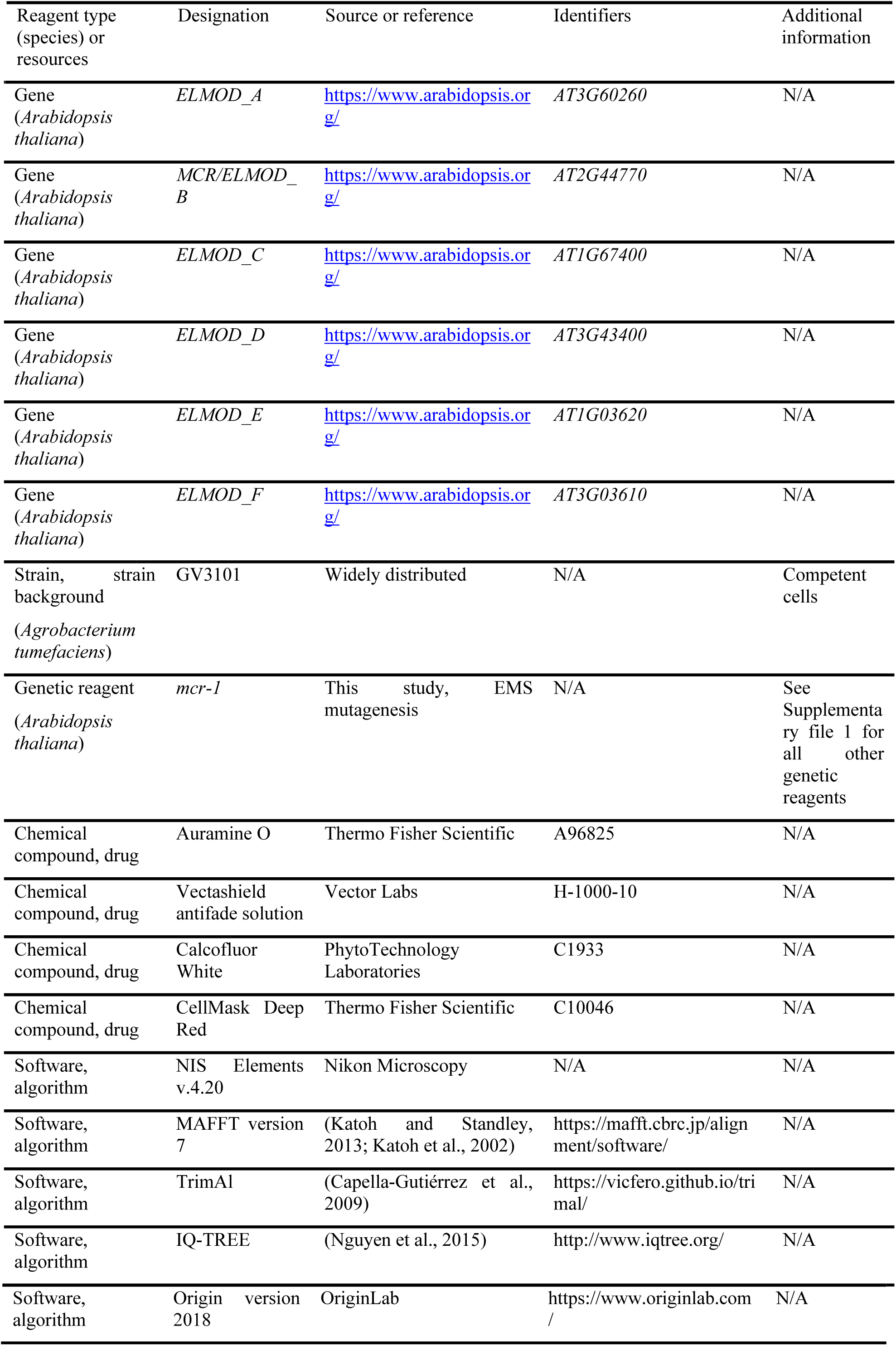

### Plant materials and growth conditions

*Arabidopsis thaliana* genotypes used in this study were either in Columbia (Col) or Landsberg *erecta* (L*er*) background. Pollen from wild-type Col-0 and L*er* has indistinguishable aperture phenotypes. The following genotypes were also used: *mcr-1*, *mcr-2*, *mcr-3*, *mcr-4*, *mcr-5* (CS853233), *mcr-6* (SALK_205528C), *mcr-7* (SALK_203827C), *elmod_c* (SALK_076565), *elmod_d* (SALK_031512), *elmod_e* (SALK_082496), *elmod_f* (SALK_010379), *inp1-1* (Dobritsa and Coerper, 2012), *inp2-1* (Lee et al., 2021), *d6pkl3-2* (Lee et al., 2018), *inp1-1 DMC1pr:INP1-YFP* (Dobritsa et al., 2018), *d6pkl3-2 D6PKL3pr:D6PKL3-YFP* (Lee et al., 2018), *tes* (SALK _113909), *MiMe* (d’Erfurth et al., 2009), *cenh3-1 GFP-tailswap* (CS66982). *mcr-1* through *mcr-4* mutants were discovered in a forward genetic screen performed on an EMS-mutagenized L*er* population (Plourde et al., 2019). *mcr-5* through *mcr-7* mutants and *elmod_c* through *elmod_f* mutants were ordered from the Arabidopsis Biological Resource Center (ABRC). Plants were grown at 20–22°C with the 16-hour light/8-hour dark cycle in growth chambers or in a greenhouse at the Biotechnology support facility at OSU.

To generate the 2n *mcr tes* plants, *mcr-1* mutant was crossed with heterozygous *tes*, double heterozygotes were recovered in F_1_ by genotyping (primers listed in Supplementary file 1), and double homozygotes were identified in F_2_ population. The generation of haploid *mcr MiMe* plants was similar to the procedure previously described (Reeder et al., 2016). In brief, *mcr-1* mutant was first crossed with plants that were triple heterozygotes for *atrec8-3*, *osd1-3*, and *atspo11-1-3* (*MiMe* heterozygotes), then the quadruple heterozygotes were identified among the F_1_ progeny by genotyping and crossed as males with *cenh3-1 GFP-tailswap* homozygous plants that were used as haploidy inducers (Ravi and Chan, 2010). 1n F_1_ progeny of this cross were identified by their distinctive morphology as described (Ravi and Chan, 2010; Reeder et al., 2016), and the triple 1n *MiMe* and quadruple 1n *mcr MiMe* mutants were identified by genotyping (primers listed in Supplementary file 1). Unlike other 1n genotypes generated by this cross, which were sterile, the 1n plants with *MiMe* mutations were fertile and produced 1n pollen via mitosis-like division and dyad formation.

### Mapping of the *MCR* locus

*mcr-1* mutant with L*er* background was crossed with Col-0, and individual F_2_ plants were screened under a dissecting microscope for the presence of the distinctive angular mutant phenotype in their dry pollen. In total, 369 plants with mutant phenotype were selected, and their genomic DNA was isolated. To map the *MCR* locus, we first conducted bulked segregant analysis, followed by the map-based positional cloning (Lukowitz et al., 2000). The insertion-deletion (InDel) molecular markers were developed based on the combined information from the 1001 Genomes Project database (1001 Genomes Consortium, 2016) and the Arabidopsis Mapping Platform (Hou et al., 2010). The *MCR* locus was mapped to a 77-kb region between markers 2-18.39 Mb (18,395,427 bp) and 2-18.47 Mb (18,472,092 bp) on chromosome 2. Molecular markers used for mapping are listed in Supplementary file 1. Out of the 25 genes located in this interval we sequenced 11 genes, prioritized based on their predicted expression patterns and gene ontology, and found that one of them, *At2g44770*, contained a missense mutation. Sequencing of the other three non-complementing EMS alleles identified in the forward genetic screen (*mcr-2* to *mcr-4*) also revealed presence of mutations in *At2g44770*.

### Inactivation of *ELMOD_A* and *ELMOD_E* with CRISPR/Cas9

Two guide RNAs against target sequences at the beginning of the *ELMOD_A* and *ELMOD_E* CDS were selected with the help of the CRISPR-PLANT platform (https://www.genome.arizona.edu/crispr/) (Xie et al., 2014) and individually cloned into the *Bsa*I site of the pHEE401E vector (Wang et al., 2015) as described (Xing et al., 2014), using, respectively, two sets of complementary primers: *elmod_a* sgRNA-F/R and *elmod_e* sgRNA-F/R (Supplementary file 1). The resulting constructs were separately transformed into the *Agrobacterium tumefaciens* strain GV3101, and then used to transform Arabidopsis Col-0 plants or *mcr-1* mutants (the latter only with the anti-*ELMOD_E* construct) using the floral-dip method (Clough and Bent, 1998). The T_1_ transformants were selected on ½ strength MS plates supplemented with 1% (w/v) sucrose, 0.8% (w/v) agar, and 50 µg/mL hygromycin, their DNA was extracted, and the regions surrounding the target sequences were sequenced. For *ELMOD_A*, five of 25 T_1_ plants had homozygous, biallelic, or heterozygous mutations. Sequencing the progeny of these plants demonstrated that all homozygous/biallelic mutants developed frame shifts in the *ELMOD_A* CDS after the codon 64 (by acquiring either a one-nt insertion three nucleotides before PAM or a one-nt deletion two nucleotides before PAM). An *elmod_a* mutant with a single A insertion, as shown in Figure 4A, and still carrying CRISPR/Cas9 transgene, was crossed with the *mcr-1* mutant to obtain the *mcr elmod_a* double mutant. For *ELMOD_E*, one out of 12 and one out of 20 T_1_ plants had biallelic mutations, respectively, in Col-0 and *mcr-1* backgrounds. In T_2_ generation, homozygous mutants with a frame shift in the CDS were identified: in *elmod_e^CR^*, a 13-nt region located four nucleotides before PAM was deleted and replaced with a different 9-nt sequence; in *mcr elmod_e^CR^*, a single A was inserted four nucleotides before PAM. These mutants were used to observe the aperture phenotypes.

### Generation of transgenic constructs and plant transformation

A 3,076-bp fragment upstream of the start codon of *MCR* was used as the *MCR* promoter for all *MCRpr* constructs. To generate the *MCRpr:gMCR* construct, the promoter and the 2,868-bp genomic fragment from the *MCR* start codon to 798 bp downstream of the stop codon were separately amplified from Col-0 genomic DNA and cloned into *Sac*I/*Nco*I sites in the pGR111 binary vector (Dobritsa et al., 2010) through In-Fusion cloning (Takara). An *Age*I site was introduced in front of the *MCR* start codon for ease of subsequent cloning. For *MCRpr:MCR CDS*, the genomic fragment was replaced with the *MCR* coding sequence, which was amplified from the *MCR* cDNA construct CD257409 obtained from ABRC. For *MCRpr:gMCR-YFP* construct, the genomic fragment of *MCR* was amplified without the stop codon and cloned upstream of *YFP* into the pGR111 binary vector (Dobritsa et al., 2010). Additionally, a 497-bp 3’ UTR region from *MCR* was then cloned downstream of *YFP*. Since we achieved phenotypic rescue and observed strong YFP signal with this construct, we used this combination of regulatory elements in all subsequent constructs for which we wanted to achieve the *MCR*-like expression. The constructs *MCRpr:gELMOD_A/C/D/E/F-YFP* were created in a similar way.

For all *EApr* constructs, a 2,163-bp fragment upstream of the start codon of *ELMOD_A* was amplified from Col-0 genomic DNA and used as the *ELMOD_A* promoter. For *EApr:gELMOD_A*, a 2,833-bp fragment, which included a 296-bp region downstream of the stop codon, was subcloned into pGR111 downstream of *EApr*. A *Bam*HI site was introduced in front of the start codon for ease of subsequent cloning. For *EApr:gELMOD_A-YFP*, a 2,534-bp genomic fragment (from the *ELMOD_A* start codon to immediately upstream of the stop codon) was cloned between the *EApr* and *YFP*. For *ELMOD_Epr:ELMOD_E-YFP*, a 1,469-bp fragment upstream of the start codon of *ELMOD_E* was amplified from Col-0 genomic DNA and used as the *ELMOD_E* promoter to replace the *MCR* promoter in *MCRpr:gELMOD_E-YFP*.

To generate the reporter constructs *MCRpr:H2B-RFP* and *EApr:H2B-RFP*, the *H2B-RFP* fusion gene was cloned into the *Bam*HI/*Spe*I sites downstream of the respective promoters in pGR111. To create constructs with single and double nucleotide substitutions, PCR-based site-directed mutagenesis was performed with IVA mutagenesis (García-Nafría et al., 2016) using *gMCR*- pGEM-T Easy, *gELMOD_A*-pGEM-T Easy and *gELMOD_E*-pGEM-T Easy as templates. The mutated sequences then replaced the respective wild-type sequences in *MCRpr:gMCR-YFP*- pGR111, *EApr:ELMOD_A-YFP*-pGR111, and *MCRpr:gELMOD_E-YFP*-pGR111. All primers used for creating constructs are listed in Supplementary file 1. All constructs were verified by sequencing and transformed by electroporation into the Agrobacterium strain GV3101 together with the helper plasmid pSoup. Agrobacterium cultures confirmed to contain the constructs of interest were then transformed into *mcr* or *mcr elmod_a* by floral dip (*mcr elmod_a* was verified to lack the anti-ELMOD_A CRISPR/Cas9 transgene).

### Confocal microscopy and image analysis

Preparation and imaging of mature pollen grains, MMC, tetrads, and free microspores were performed as previously described (Reeder et al., 2016). Imaging was done on a Nikon A1+ confocal microscope with a 100× oil-immersion objective (NA = 1.4), using 1× confocal zoom for anthers, 3× zoom for pollen grains, 5× zoom for MMC and tetrads, and 5× or 8× zoom for free microspores. For imaging mature pollen grains, pollen was placed into a ∼10-μL drop of auramine O working solution (0.001%; diluted in water from the 0.1% (w/v) stock prepared in 50 mM Tris-HCl), allowed to hydrate for ∼5 min, covered with a #1.5 cover slip, and sealed with nail polish. Exine was excited with a 488-nm laser and fluorescence was collected at 500-550 nm. To count aperture number, images from the front and back view of pollen grain were taken. If some apertures were present on sides of a pollen grain not directly visible by focusing on the front and on the back, then z-stacks were taken (step size = 500 nm) and 3D images were reconstructed using NIS Elements software v.4.20 (Nikon) and used for counting aperture number.

For imaging cells of the developing pollen lineage, anthers were dissected out of stage-9 flower buds and placed into a small drop of Vectashield antifade solution supplemented with 0.02% Calcofluor White and 5 μg/mL membrane stain CellMask Deep Red. Cells in the pollen lineage were released by applying gentle pressure to the coverslip placed over the anthers. To obtain fluorescence signals, the following excitation/emission settings were used: RFP, 561 nm/580-630nm; YFP, 514 nm/522-555 nm; Calcofluor White, 405 nm/424-475 nm; CellMask Deep Red, 640 nm/663-738 nm. Z-stacks of tetrads were obtained with a step size of 500 nm and 3D reconstructed using NIS Elements v.4.20 (Nikon).

To compare the YFP fluorescence intensity in three different lines of *mcr MCRpr:gMCR*-*YFP* or *mcr elmod_a EApr:gELMOD_A*-*YFP*, tetrads were prepared simultaneously and imaged on the same day under identical acquisition settings on Nikon A1+ confocal microscope. The mean YFP signal intensities in nucleoplasm and cytoplasm of tetrads (n ≥ 15) were separately measured with the help of the ROI (Region of interest) statistics function in NIS Elements v.4.20 (Nikon). For each tetrad, a single optical section showing both nucleoplasm and cytoplasm was selected and analyzed.

### Sequence retrieval and phylogenetic analysis of the plant ELMOD family

ELMOD family members in Arabidopsis have the following accession numbers: *ELMOD_A*, *At3g60260*; *MCR*, *At2g44770*; *ELMOD_C*, *At1g67400*; *ELMOD_D*, *At3g43400*; *ELMOD_E*, *At1g03620*; *ELMOD_F, At3g03610*. The phylogenetic tree of Arabidopsis ELMOD proteins in Figure 3E was built using the neighbor-joining (NJ) algorithm of MEGA X (Kumar et al., 2018), with bootstrap support calculated for 1,000 replicates.

Sequences of ELMOD proteins from species across the plant kingdom were retrieved from the Phytozome v.12 database (https://phytozome.jgi.doe.gov/pz/portal.html) and the 1,000 Plants (1KP) database (https://db.cngb.org/onekp/, last accessed in May 2020) (Wickett et al., 2014). MCR protein sequence was used as a query for an online BLASTP search of these databases with default parameters. The protein sequences with the E-value ≤1e-10, sequence identity ≥30%, and Bit-Score ≥60 were identified as ELMODs and further confirmed by a local BLASTP search using each of the other Arabidopsis ELMODs as a query. In the cases when two or more proteins were potentially translated from the same gene, the one providing the best match with the query was selected. In total, 561 ELMOD protein sequences from 178 representative species belonging to eudicots (36 species/195 sequences), monocots (14/94), magnoliids (20/64), basal angiosperms (5/13), gymnosperms (17/62), ferns (17/44), lycophytes (20/29), bryophytes (37/47; including 18 sequences from 15 liverworts, 6 sequences from 5 hornworts, and 23 sequences from 17 mosses)), and green algae (12/13) were retrieved and used for phylogenetic analysis.

Multiple sequence alignment was performed using MAFFT v7.017 (Katoh and Standley, 2013; Katoh et al., 2002) with the L-INS-i algorithm and default parameters. Sites with greater than 20% gaps were trimmed by TrimAl (Capella-Gutiérrez et al., 2009) and manually inspected for overhangs. ModelFinder (Kalyaanamoorthy et al., 2017) (accessed through IQ-TREE (Nguyen et al., 2015)) was run to find the best-fit amino acid substitution model. The alignment in Figure 3—figure supplement 1 was visualized with Espript3.0 (Gouet et al., 1999). Phylogenetic trees were constructed using IQ-TREE with the Maximum Likelihood (ML) method, SH-aLRT test, and ultrafast bootstrap with 1,000 replicates. For the tree on Figure 8A, containing sequences from across the plant kingdom, 267 sequences were used, including all sequences retrieved from green algae, bryophytes, lycophytes, ferns, gymnosperms, and basal angiosperms, as well as 24 sequences from magnoliids, 19 sequences from 3 monocots, and 16 sequences from 3 eudicots. For the tree on Figure 8—figure supplement 1, containing only angiosperm sequences, we used all 366 sequences retrieved for this group. Phylogenetic trees were visualized in iTOL v.5 (Letunic and Bork, 2021) and can be accessed at http://itol.embl.de/shared/Zhou3117.

### Expression pattern analysis of the Arabidopsis *ELMOD* genes

RNA-seq data for different tissues/developmental stages of six Arabidopsis *ELMOD* genes were obtained from the TRAVA database (http://travadb.org/) (Klepikova et al., 2016). The ‘Raw Norm’ option was chosen for read counts, and default settings were used for all other options. The retrieved RNA-seq data were presented as a bubble heatmap using TBtools (Chen et al., 2020).

## QUANTIFICATION AND STATISTICAL ANALYSIS

Quantification of aperture numbers and YFP signal was done with NIS Elements v.4.20 software (Nikon). For each line, the aperture number of 160 pollen grains from at least three different plants were counted and the mean YFP fluorescence of at least 15 tetrads from the same plants was measured. Graphs were generated using Microsoft Excel or Origin version 2018 software. Binary comparisons were performed using a two-tailed Student’s t-test in Microsoft Excel; results with the *p* values below 0.05 were judged significantly different. The *p* values are represented as *** (*p* < 0.001); ** (*p* < 0.01); * (*p* < 0.05). All error bars represent standard deviation (SD). For all boxplots, the box defines the first and third quartile, the central line depicts the median, and the small square in the box represents the mean value. Whiskers extend to minimum and maximum values. Outliers are indicated as 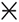. Different shapes show individual samples. Details of statistical analysis, number of quantified entities (n), and measures of dispersion can be found in the corresponding figure legends.

## Acknowledgements

Funding for this project was provided by the US National Science Foundation (MCB-1817835) (awarded to A.A.D.). We also acknowledge the support of the OSU Mayers Undergraduate Summer Research Scholarship to P.A., the NSF-REU supplement funding to P.A. and A.H., and the iCAPS internship from the OSU Center for Applied Sciences to A.H.

## Author contributions

Y.Z. and A.A.D. conceived and designed the experiments. Y.Z., P.A., S.H.R., B.H.L., A.H., and A.A.D. performed the experiments. Y.Z. and A.A.D. analyzed the data and wrote the manuscript.

## Additional files

Supplementary files

• Supplementary file 1. Primers, molecular markers, and mutants/transgenic lines used in this study.

## Data availability

The authors declare that all data supporting the findings of this study are included in the paper and the supplementary files.

---

**Figure 1—figure supplement 1.**
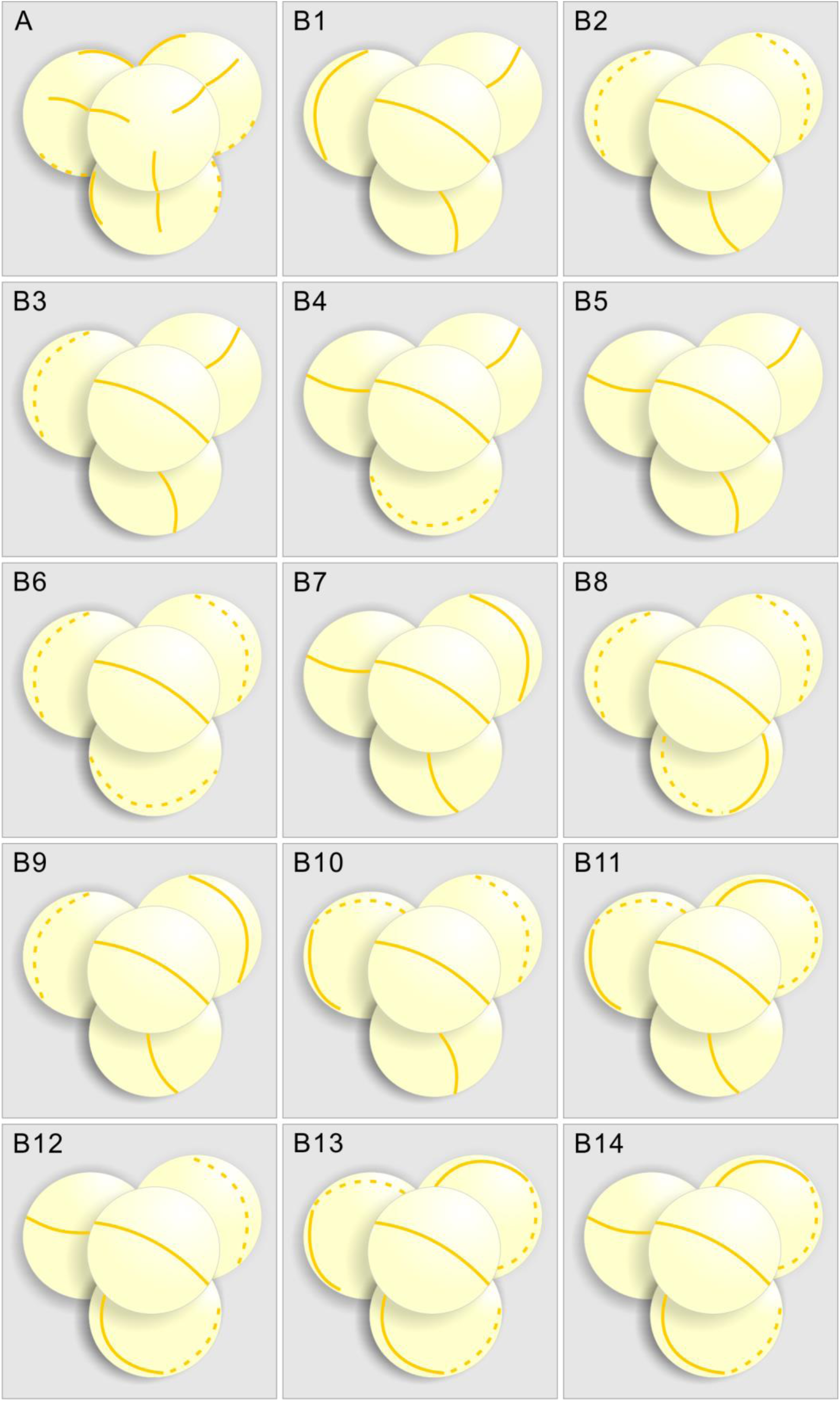
Diagrams summarizing the INP1-YFP localization in *inp1* and *mcr* tetrads, based on confocal imaging and 3-D reconstruction of *DMC1pr:INP1-YFP*-expressing tetrads. (A) Positions of three equidistant lines formed by INP1-YFP in tetrad-stage *inp1* microspores always appear coordinated between the sister microspores, with each line in one microspore facing a line in one of its sisters. (B) Examples of placement of INP1-YFP ring-shaped lines in 14 *mcr* tetrads, which suggest that the lines in sister microspores are positioned independently. In all tetrads, the INP1-YFP lines in front-facing microspores (with the polar axis perpendicular to the plane of image) were oriented the same way to compare the positioning of the lines in three sister microspores between the tetrads. Solid lines and dotted lines represent the INP1-YFP lines that are, respectively, visible and invisible in that view.

**Figure 1—figure supplement 2.**
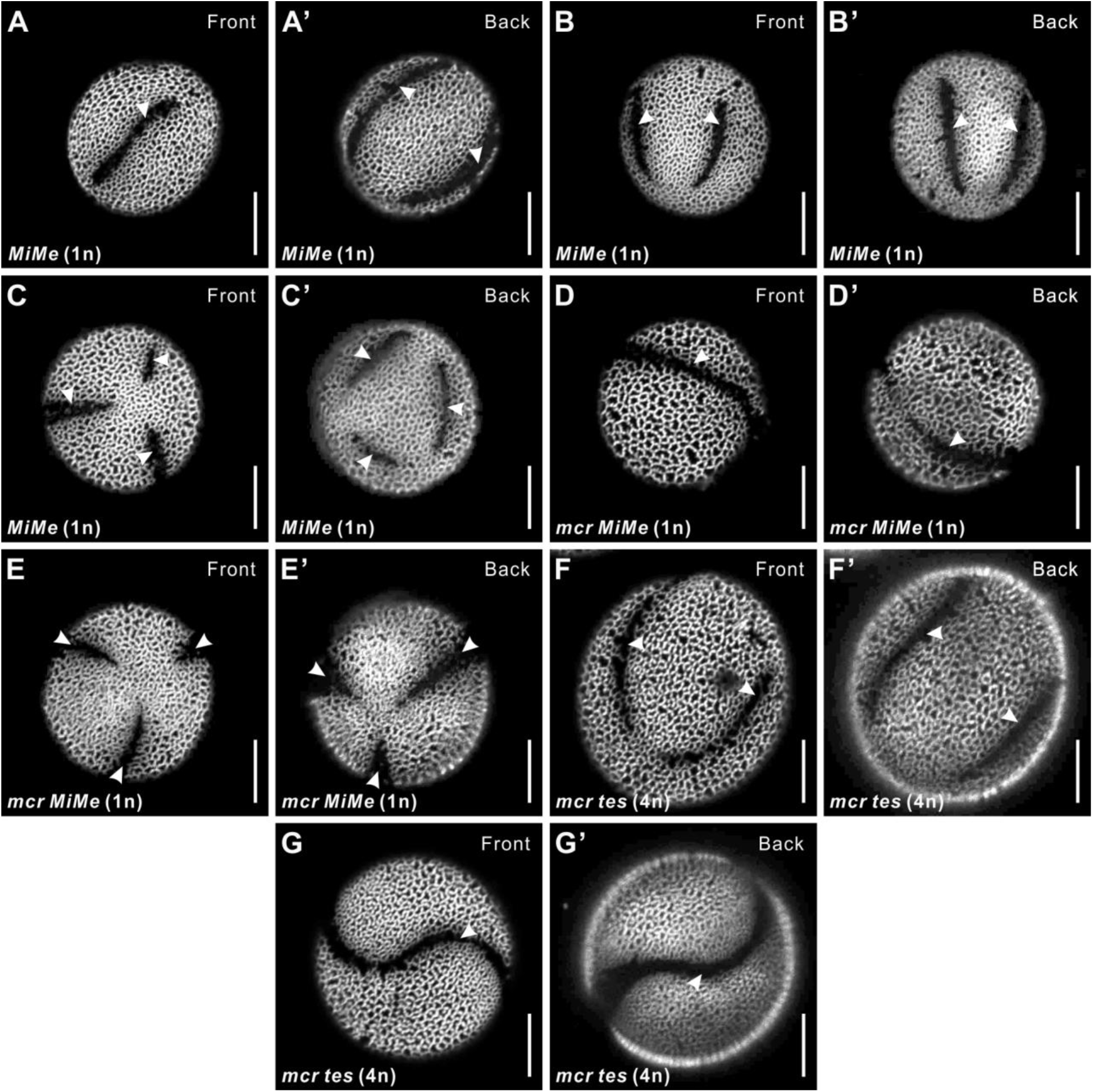
The reducing effect of *mcr* mutations on aperture number is manifested across different ploidy levels and arrangements of microspores. (A–C’) Representative images of 1n *MiMe* pollen with 3 apertures (A–A’), 4 apertures (B–B’), and 6 apertures (C–C’). (D–E’) Representative images of 1n *mcr MiMe* pollen with *mcr*-like aperture (D–D’) and 3 apertures (E–E’). (F–G’) Representative images of 4n *mcr tes* pollen with 4 apertures (F–F’) and fused apertures (G–G’). Apertures are indicated with arrowheads. Scale bars, 10 μm.

**Figure 3—figure supplement 1.**
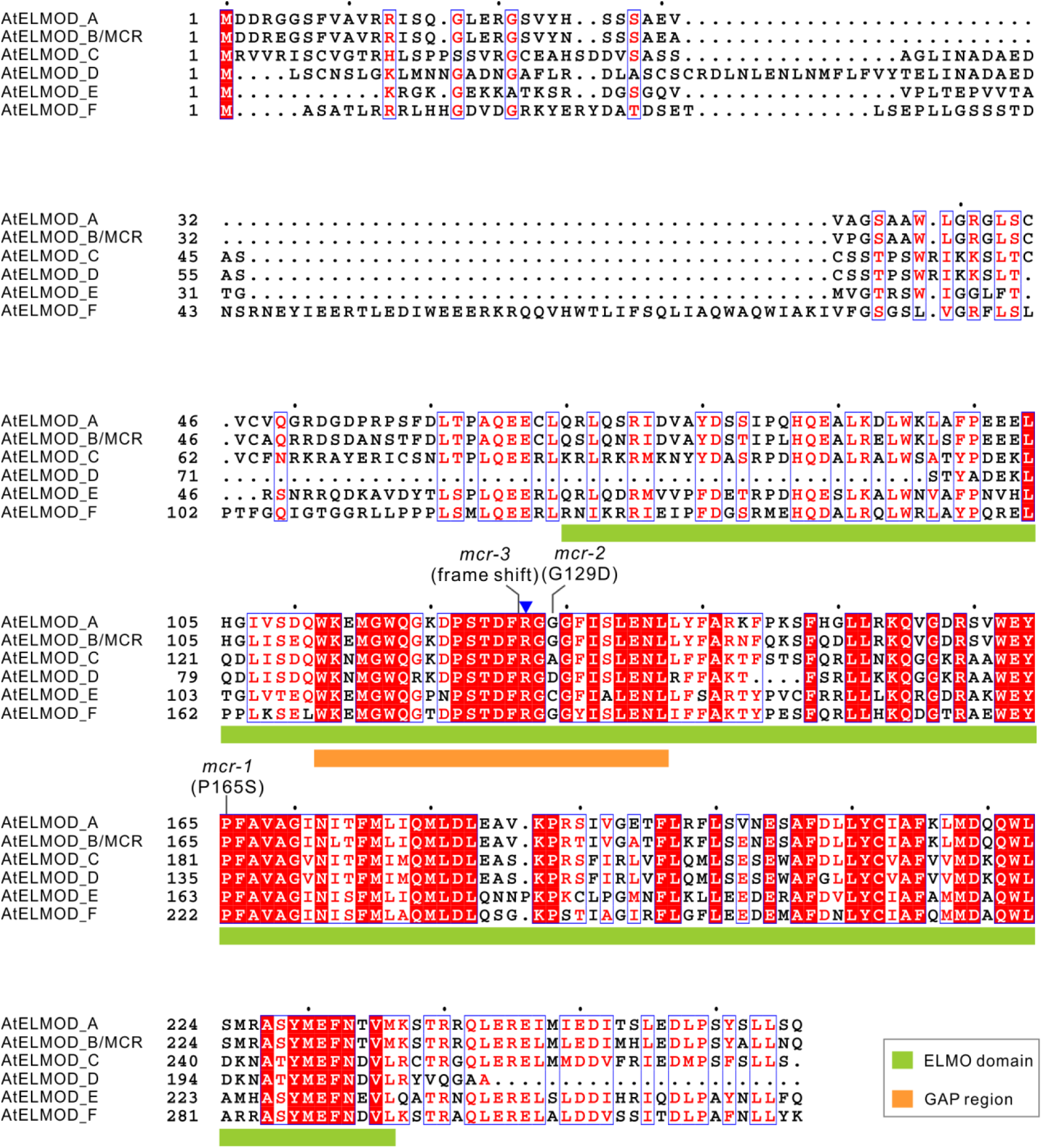
Protein sequence alignment of Arabidopsis ELMOD proteins. Multiple sequence alignment was conducted by MAFFT and visualized by Espript3.0. Positions of ELMO domains and GAP regions in these proteins are indicated with a green box and an orange box, respectively. The mutated sites of *mcr-1*, *mcr-2*, and *mcr-3* are indicated. Blue triangle indicates the highly conserved Arg (R127 of MCR and ELMOD_A) in the GAP region.

**Figure 3—figure supplement 2.**
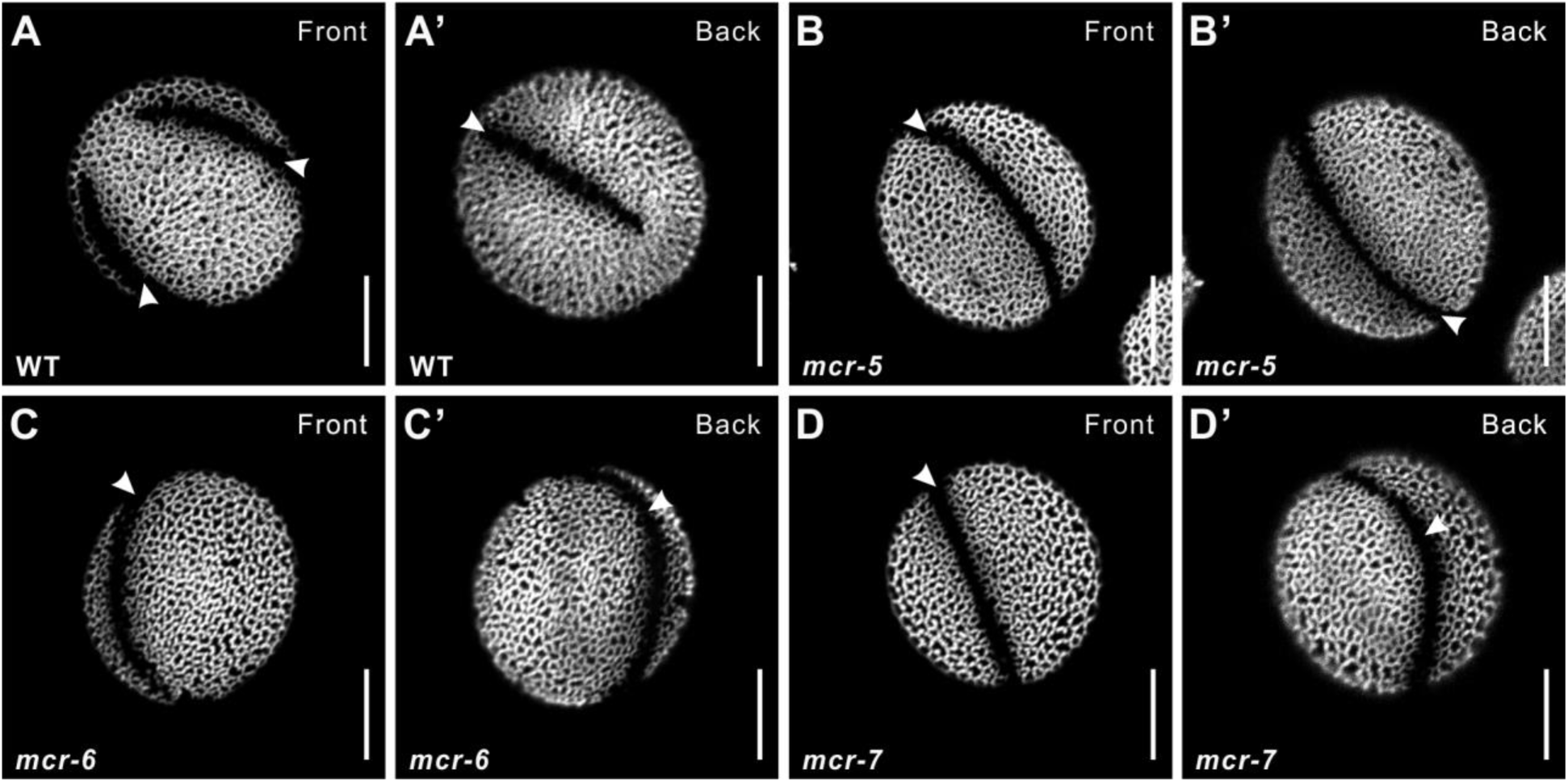
T-DNA insertion mutants of *MCR* produce pollen with a single ring-shaped aperture. Pollen from wild type (A–A’) and three T-DNA insertion alleles of *MCR* (B–D’). Apertures are indicated with arrowheads. Scale bars, 10 μm.

**Figure 7—figure supplement 1.**
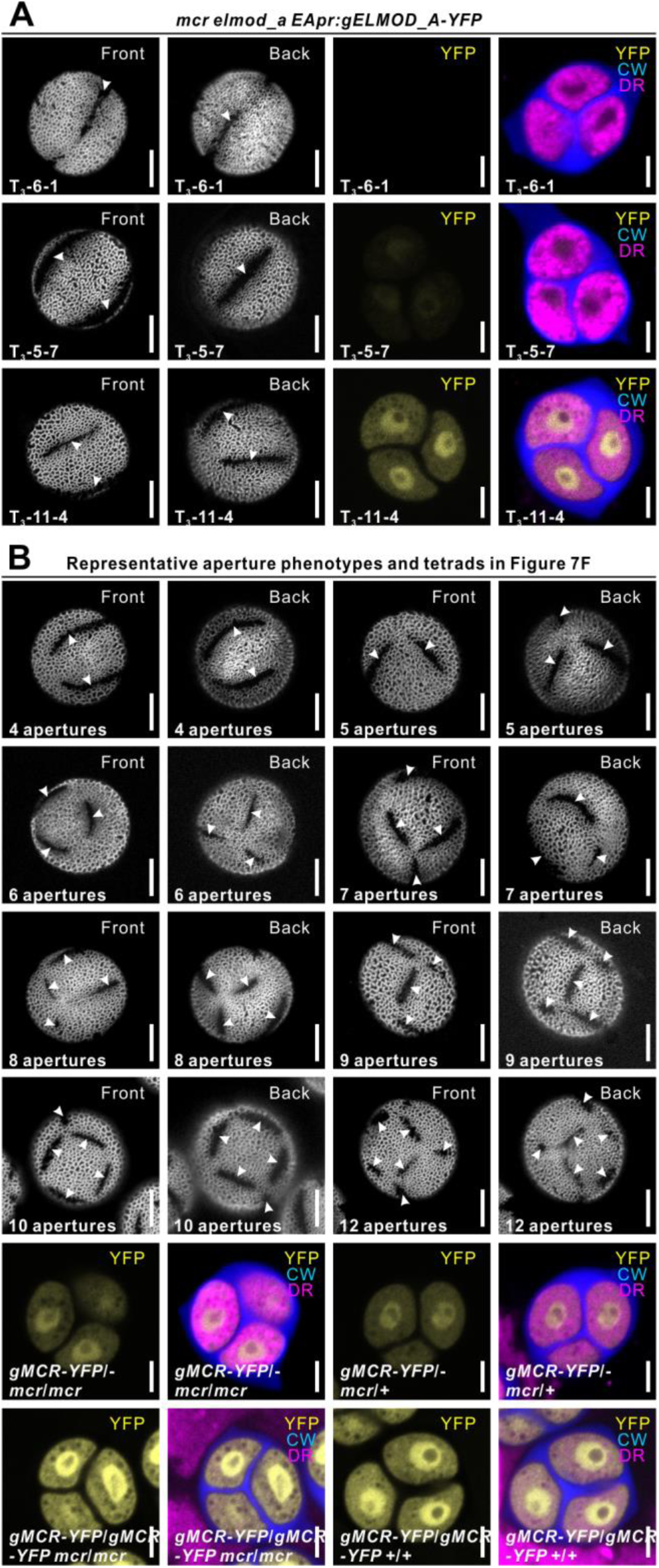
Representative aperture phenotypes and tetrads related to Figure 7. (A) Representative images of pollen grains and tetrads used in Figure 7D. (B) Representative images of pollen grains and tetrads used in Figure 7F. Apertures are indicated with arrowheads. Scale bars, 10 μm for pollen and 5 μm for tetrads.

**Figure 8—figure supplement 1.**
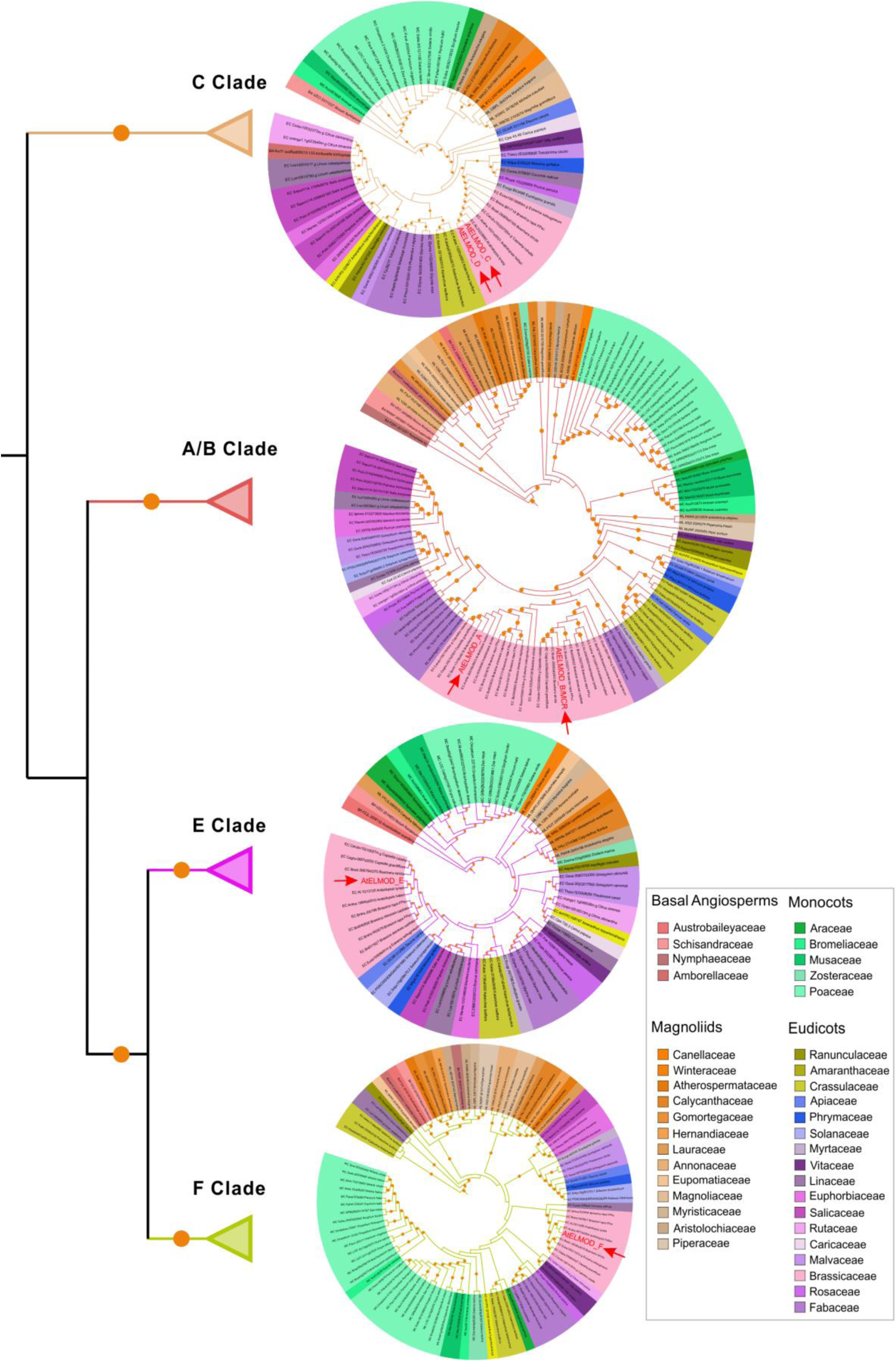
Angiosperm ELMOD proteins cluster into four clades. Maximum likelihood phylogenetic tree of ELMOD proteins from angiosperms. The four distinct clades have been collapsed and details of each clade are presented as the pruned circular tree on the right. Label color shows the taxonomic group of each protein as indicated on the right. Orange circles indicate bootstrap values higher than 70%. Red arrowheads indicate the Arabidopsis ELMOD proteins. The complete tree can be accessed at http://itol.embl.de/shared/Zhou3117.

**Figure 9—figure supplement 1.**
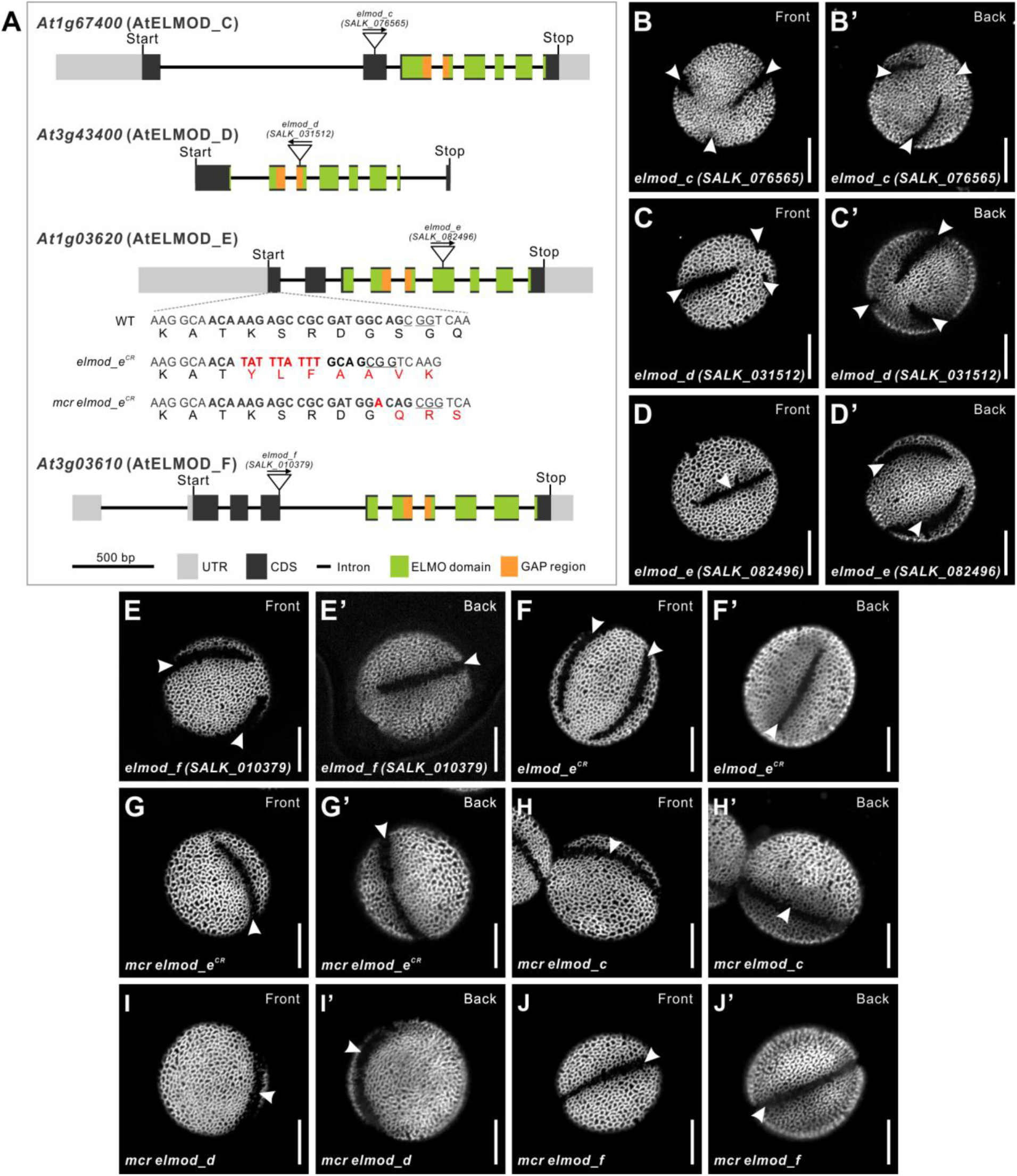
Disruptions of Arabidopsis *ELMOD_C*, *ELMOD_D*, *ELMOD_E*, and *ELMOD_F* do not affect aperture patterns. (A) Diagram of the *ELMOD_C* (*At1g67400*), *ELMOD_D* (*At3g43400*), *ELMOD_E* (*At1g03620*), and *ELMOD_F* (*At3g03610*) genes. T-DNA insertion sites are indicated for each gene. For *ELMOD_E*, CRISPR alleles (*elmod-e^CR^*) generated in the wild-type and *mcr* backgrounds are shown. The 20-bp target sequence next to the underlined protospacer adjacent motif is indicated in bold. Nucleotide and amino acid changes are indicated with red capital letters. (B–F’) Pollen grains of single T-DNA insertion mutants of *ELMOD_C*, *ELMOD_D*, *ELMOD_E*, *ELMOD_F* and the CRISPR/Cas9 mutant of *ELMOD_E* (*elmod_e^CR^*). (G–J’) Pollen grains of the double mutants *mcr elmod_e^CR^* (G–G’)*, mcr elmod_c* (H–H’)*, mcr elmod_d* (I–I’), and *mcr elmod_f* (J–J’). Apertures are indicated with arrowheads. Scale bars, 10 μm.

**Figure 9—figure supplement 2.**
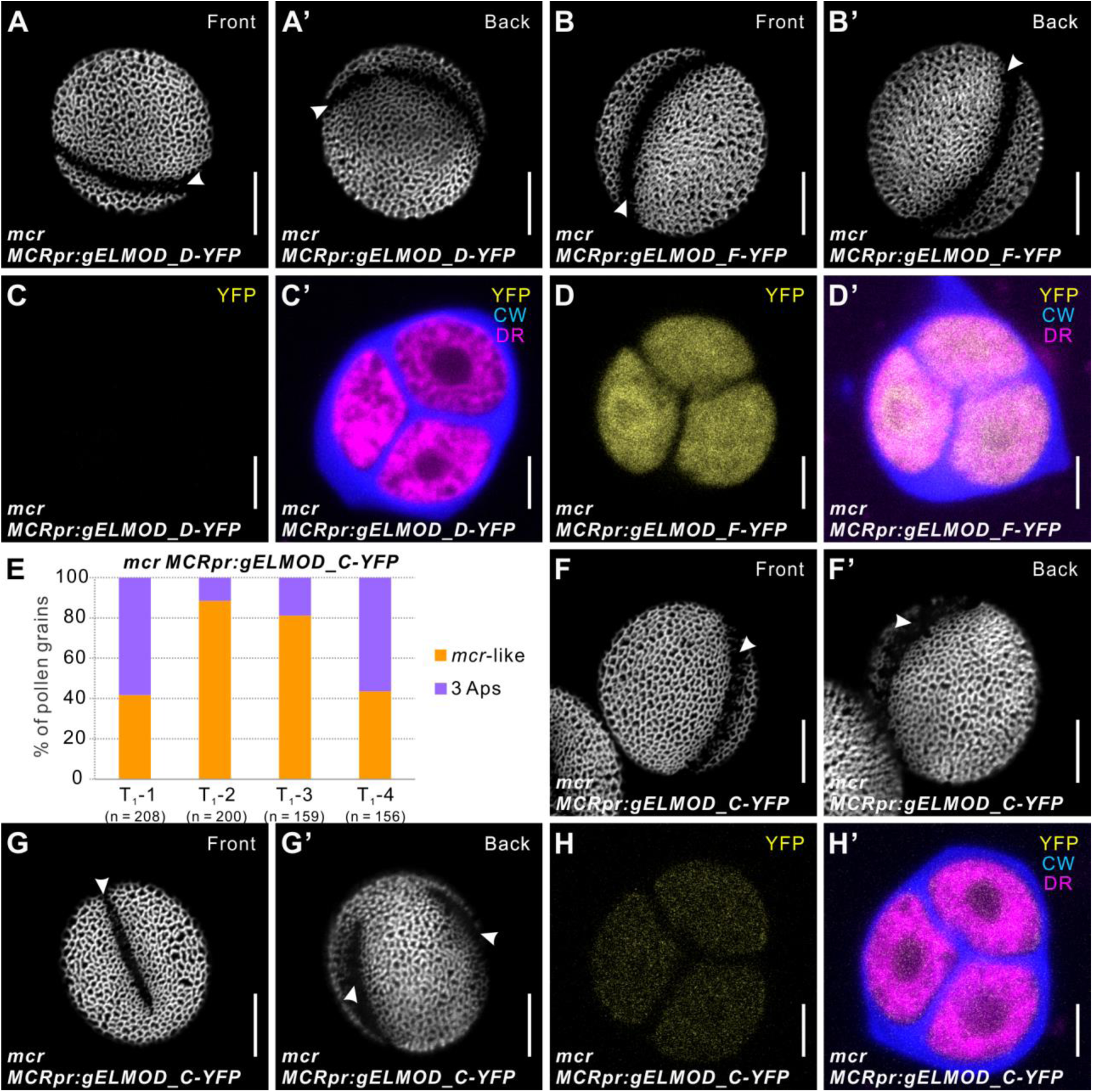
ELMOD_C, but not ELMOD_D and ELMOD_F, can partially substitute for MCR in aperture formation. (A–B’ and F–G’) Pollen grains from *mcr MCRpr:gELMOD_D-YFP* (A–A’), *mcr MCRpr:gELMOD_F-YFP* (B–B’), and *mcr MCRpr:gELMOD_C-YFP* (F-G’). (C–D’ and H–H’) Confocal images of *mcr* tetrads expressing YFP fusions of ELMOD_C, ELMOD_D, and ELMOD_F. (C–C’) There is no observable YFP fluorescence in the *mcr* tetrads expressing *MCRpr:gELMOD_D-YFP.* (D–D’) *mcr* tetrads expressing *MCRpr:gELMOD_F-YFP* show strong YFP fluorescence. (H–H’) *mcr* tetrads expressing *MCRpr:gELMOD_C-YFP* have detectable YFP fluorescence. Adjacent panels show YFP fluorescence (α) and merged fluorescent signal (α’) from YFP, Calcofluor White (CW), and CellMask Deep Red (DR). (E–E’) Percentage of pollen grains with indicated number of apertures in pollen populations from the T_1_ plants of *mcr MCRpr:gELMOD_C-YFP*. Number of analyzed pollen grains is indicated. Apertures are indicated with arrowheads. Scale bars, 10 μm for pollen and 5 μm for tetrads.

**Figure 10—figure supplement 1.**
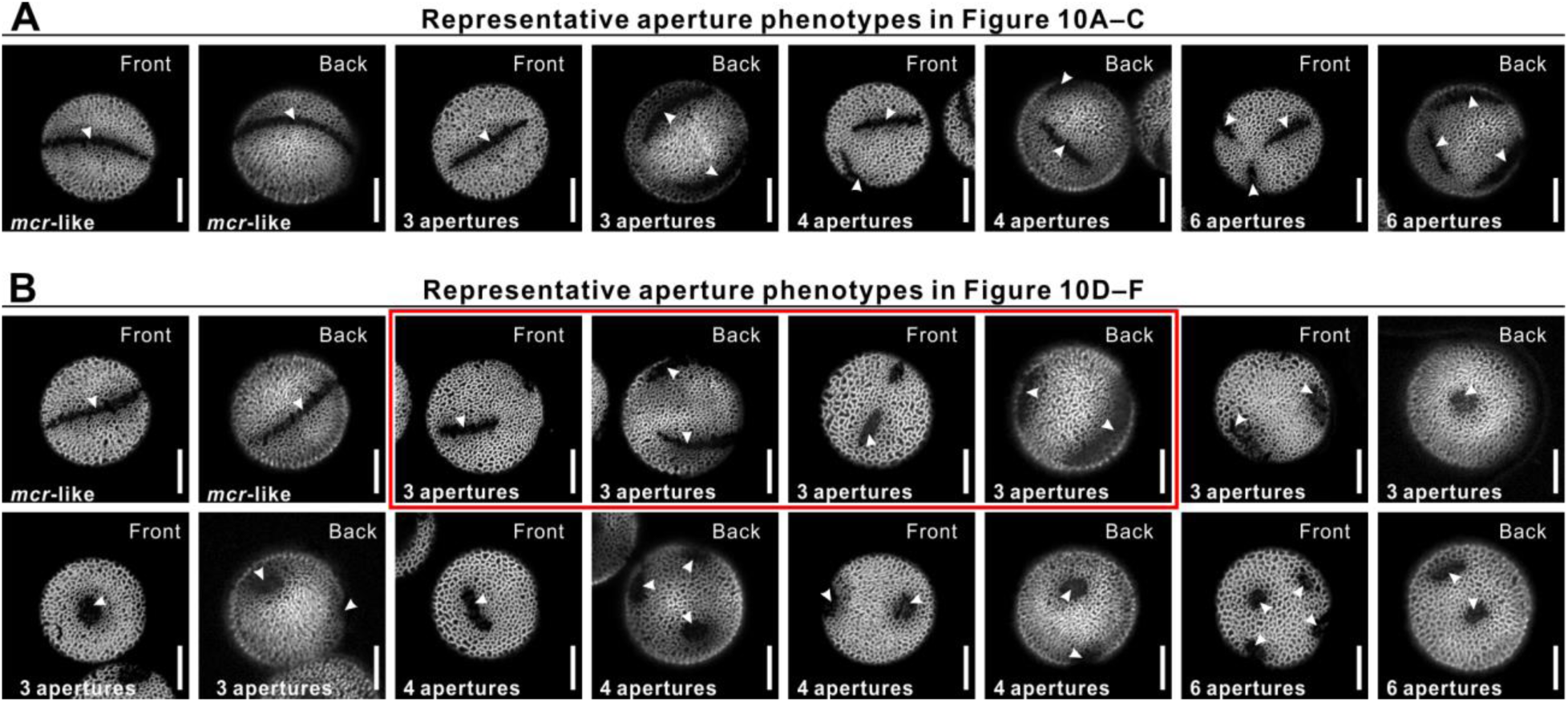
Representative aperture phenotypes observed in T_1_ plants related to Figure 10. (A) Representative aperture phenotypes (all furrows) observed in T_1_ plants related to Figure 10A–C. (B) More diverse aperture phenotypes observed in T_1_ plants related to Figure 10D–F. Red box highlights the most common aperture morphologies of three normal or, sometimes, disconnected furrows observed in the T_1_ plants related to Figure 10D. Three round apertures were only found in the T_1_ plants related to Figure 10E and Figure 10F. ≥4 apertures were mostly round. Apertures are indicated with arrowheads. Scale bars, 10 μm.

